# Plasticity and Dynamics of Hematopoietic Cells within Bone Marrow Microenvironment in Leukemia

**DOI:** 10.1101/2024.04.02.587680

**Authors:** Chuijin Wei, Shumin Xiong, Yi Zhou, Liaoliao Dong, Ping Yu, Yunhan Tang, Ren Zhou, Beiwen Ni, Jian Hou, Guang Liu, Lin Cheng

## Abstract

Extensive research has been conducted on the plasticity of malignant cells and nonmalignant cells in solid tumor. However, the plasticity of bone marrow hematopoietic cells in leukemia have remained largely unexplored. In this study, we aimed to investigate cell changings in hematopoietic cells through lineage tracing across various types of leukemias. We had compiled a landscape of leukemia and constructed phylogenetic trees of hematopoietic cells through utilizing massively parallel scRNA-seq data, mtDNA mutation and SNP analysis. Based on the observed cell changings, we identified several types of cell changings, including transdifferentiation, dedifferentiation, and state transition, except for differentiation and expansion. In AML and CMML, GMPs and neutrophils showed a higher potential for transferring to other cell types. In BPDCN, pDCs were less prone to switching to other cell types, while T cells demonstrated high plasticity. In B-ALL and B-CLL, B-ALL blast cells and B-CLL blast cells emerged at the most dynamic state. The dynamics of hematopoietic cells in AML, BPDCN and ALL changed along with the clinical process. Extrinsic factors within the leukemia microenvironment may influence the cell changings. Regulons encountered an intermediate cell state during the process of transition to myeloid cells and erythroid cells. We also found a correlation between B-common blast cells and T cells, suggesting a potential transition from B lymphoblastic leukemia to T lymphoblastic leukemia. In conclusion, our study unveiled the distinct plasticity and dynamics of hematopoietic cells in various types of leukemia. This sheds light on the possibility of targeting cell changes as a new strategy for leukemia treatment and improving current immunotherapy.

## Introduction

Leukemia is a malignant hematopoietic disease characterized by the abnormal proliferation of specific cell types, including acute myeloid leukemia (AML) (*1*), chronic myelomonocytic leukemia (CMML) (*2*), acute lymphoblastic leukemia (ALL) (*3, 4*), chronic lymphoblastic leukemia (CLL) (*5*), blastic plasmacytoid dendritic cell neoplasm (BPDCN) (*6*) and others. These leukemias involve the clonal expansion of cells, leading to poor prognosis. The bone marrow microenvironment in leukemia plays a crucial role, encompassing the heterogeneity and diverse functions of hematopoietic cells, which have been extensively researched (*7*). However, certain questions about leukemia hematopoietic cell plasticity remain unanswered. The use of single-cell RNA sequencing (scRNA-seq) technique, which provides more previously unnoticed information about hematopoietic cells and enables the identification of more detailed subtypes with transcriptional and genetic profiles, is a valuable tool in addressing these unresolved questions.

The bone marrow serves as the primary site for adult hematopoiesis, wherein mature hematopoietic cells are generated from hematopoietic stem and progenitor cells. This is widely recognized as a path of multi-layered, multi-branched, top-down differentiation. However, this canonical perception is gradually being challenged by the recent identification of hematopoietic cell reprogramming both *in vivo* and *in vitro* (*8*). Notably, mature hematopoietic cells can undergo transdifferentiation from one type to another or dedifferentiation into immature progenitors or stem cells *in vitro*, achieved through the overexpression of exogenous transcription factors or exposure to chemical compounds and cytokines (*9–14*). In the context of leukemia, reports indicate instances such as T-lymphoblastic lymphoma transdifferentiating into mature myeloid hematopoietic neoplasms (*15*) and CAR-T cells exhibiting a B-cell immunophenotype in B-ALL (*16*). In addition, under conditions of hypoxia, B lymphoma cells are capable of converting to erythroblast-like cells (*17*). Given the heterogeneity of bone marrow hematopoietic cells and the complex nature of their microenvironment in leukemia, the question of whether other hematopoietic cells undergo dynamic state transitions or fate changes remains largely unexplored.

In contemporary research, various lineage tracing methods have been developed to track cell changings both *in vivo* and *in vitro*. Exogenous barcodes are introduced in model animals and cell experiments to monitor these changes (*18, 19*). However, the use of exogenous barcodes in human samples presents a limitation. To address this, we employed natural mitochondrial DNA (mtDNA) mutations and chromosome single nucleotide polymorphism (SNP), coupled with transcriptional profiles, which are all simultaneously derived from massive scRNA-seq data, to construct phylogenetic trees of hematopoietic cells within the bone marrow microenvironment, unraveling the dynamics of hematopoietic cells (Fig. 1A). Utilizing an atlas of leukemia cells, we created a comprehensive landscape that mapped the dynamics of hematopoietic cells in the bone marrow microenvironment. Through these mappings, we identified cell changings in leukemia hematopoietic cells along with the clinical process. Our analysis further focused on extrinsic factors and regulons as undermining mechanisms that govern the dynamics of hematopoietic cells. Additionally, we found that B lymphoblastic leukemia has the potential to transition to T lymphoblastic leukemia. This study provides a new perspective and systematic view on the dynamics of hematopoietic cells within the leukemia bone marrow microenvironment. It also improves our understanding of the heterogeneity and plasticity of hematopoietic cells. Furthermore, it holds the potential to improve the efficiency of current clinical immunotherapy.

**Figure 1.**
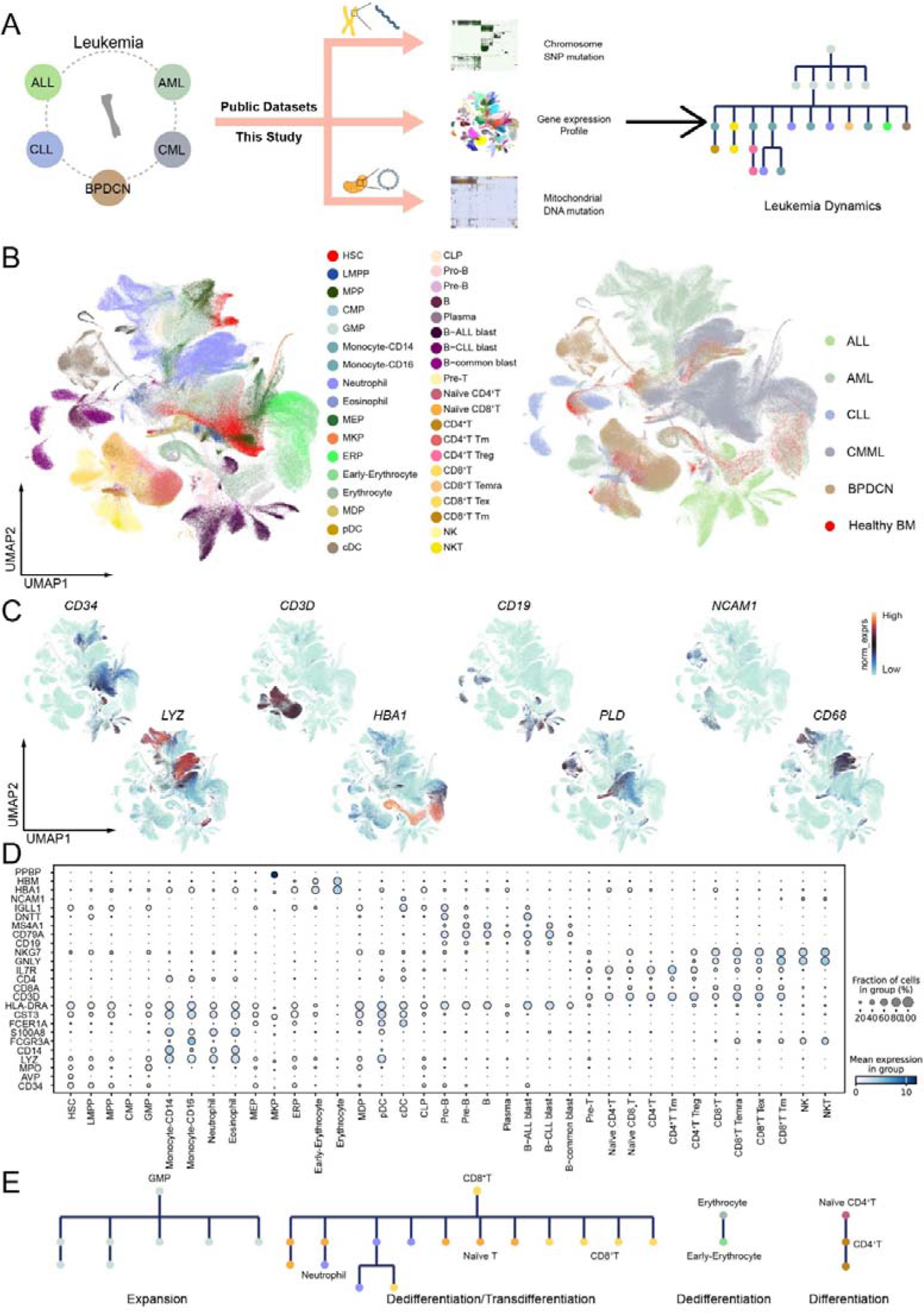
Identification of bone marrow hematopoietic cells in various leukemia types. (A) Schematic of the workflow for lineage tracing and dynamics analysis on leukemia microenvironment using single-cell transcriptome, mtDNA mutation and chromosome SNP mutation of hematopoietic cells. (B) UMAP plot showing the major lineages (left) and origins (right) of hematopoietic cells in leukemia and healthy bone marrow samples. (C) UMAP plots showing the marker genes expression for the major lineages of hematopoietic cells. (D) Bubble heatmap showing expression levels of selected signature genes in hematopoietic cells. Dot size indicates fraction of expressing cells, while color bases on mean expression levels. (E) Phylogenetic trees showing cell changings of the bone marrow hematopoietic cells in leukemia containing expansion, differentiation, dedifferentiation and transdifferentiation

## Results

### Construction of Atlas of Bone Marrow Hematopoietic Cell Lineages Based on Single Cell Transcriptional Landscape and Phylogenetic Trees

To construct a comprehensive transcriptional landscape of leukemia cells, scRNA-seq data were predominantly obtained from bone marrow hematopoietic cells in approximately 100 samples, encompassing more than 50 patients diagnosed with one of the five leukemias. This dataset included newly sequenced data from B-ALL and B-CLL (Fig. S1A, Table S1). Following rigorous quality control and filtration procedures, we compiled a dataset comprising over 500,000 hematopoietic cells derived from these leukemia samples. Additionally, healthy bone marrow hematopoietic cells were collected as a reference to delineate subtypes and cell states (Fig. 1B).

To characterize hematopoietic cell subtypes and mitigate batch effects across various datasets, we utilized the Scanpy (*20*) integrating function for analysis. Unsupervised graph-based clustering was conducted on hematopoietic cells, leading to the identification of four major lineages: myeloid cells, lymphoid cells, erythroid cells, and progenitor cells, based on canonical cell markers. Specific high expressions of *CD34*, *CD3D*, *CD19*, *NCAM1*, *LYZ*, *HBA1*, *PLD* and *CD68* characterized myeloid cells, lymphoid cells, erythroid cells, and progenitor cells, respectively (Fig. 1C). Myeloid cells were further categorized into three subsets, including monocytes (CD14^+^ monocytes and CD16^+^ monocytes), neutrophils and eosinophils. Dendritic cells (DCs) exhibited two distinct subsets, namely conventional DCs (cDCs) and plasmacytoid DCs (pDCs) (*21*). Lymphoid cells encompassed T cells, B cells, and NK cells, with CD4^+^ T cells and CD8^+^ T cells playing crucial roles in immune responses. Distinctions in immune function led to the identification of CD4^+^ memory T cells (CD4^+^ T_m_), CD4^+^ regular T cells (CD4^+^ T_reg_), CD8^+^ T terminally differentiated effector memory or effector cells (CD8^+^ T_emra_), and CD8^+^ exhausted T cells (CD8^+^ T_ex_), distinguishable by *NKG7*, *GNLY*, *IL7R* and *GZMK*, respectively. Recent research had highlighted that exhausted T cells exhibited decreased immune function and contribute to tumor progression (*22*). In B-ALL and B-CLL datasets, numerous B cells, distinct from normal B cells, were identified as B-ALL blast cells, B-CLL blast cells and B-common blast cells according to their source. Erythrocyte progenitor cells (ERPs), early-erythrocytes and mature erythrocytes were differentiated based on varying expression levels of *HBA1* and *HBM*. Stem and progenitor cells included hematopoietic stem cells (HSCs), multipotent progenitors (MPPs), lymphoid-primed multipotent progenitors (LMPPs), common myeloid progenitors (CMPs), megakaryocyte/erythrocyte progenitors (MEPs), megakaryocyte progenitors (MKPs), granulocyte/monocyte progenitors (GMPs), and common lymphoid progenitors (CLPs), expressing specific markers (*CD34*, *AVP, MPO* and *PPBP*) (Fig. 1D). In AML and CMML datasets, GMPs and neutrophils constituted a significant proportion. B cells predominated in the B-ALL and B-CLL datasets, while pDC cells were the most abundant in the BPDCN datasets (Fig. S1B).

Mitochondria played a pivotal role in cellular metabolism and are distinctive organelles possessing their own genome. Notably, mtDNA mutation could be captured through scRNA-seq (Fig. S1C). Previous studies had leveraged mtDNA mutations to establish clonal relationships among cells (*23–25*). However, relying solely on mtDNA mutations can lead to false positives that may be due to mitochondrial transfer between cells. To address this issue, we incorporated SNPs that were also inferred from the scRNA-seq data (Fig. S1D). The derived-SNPs data can also contribute to constructing clonal relationships among cells (*26, 27*). Nevertheless, our prior research demonstrated that this approach lacks precision in tracing cell changes within the microenvironment. To enhance lineage tracing accuracy, we integrated the two endogenous genetic codes and identified cell changings with a high correlation in phylogenetic trees (Fig. S1E), similarly to our previous report on constructing phylogenetic trees of immune cells within solid tumors (*28*). Based on the phylogenetic trees and the transcriptional landscape, atlas of bone marrow hematopoietic cell lineages was constructed (Fig. 1E). According to the atlas, bone marrow hematopoietic cells in leukemia underwent expansion, differentiation, dedifferentiation, state transition and transdifferentiation at different frequencies.

### Dynamics of Hematopoietic Cells Along with Clinical Progress in AML

To construct a comprehensive transcriptional landscape of AML hematopoietic cells, scRNA-seq data from 10 patients, spanning the clinical progress of AML from diagnosis to minimal residual disease (MRD) samples and relapse, were integrated. The landscape revealed the identification of 18 distinct cell types, including HSCs, MPPs, LMPPs, CMPs, GMPs, monocytes, neutrophils, MEPs, ERPs, early-erythrocytes, erythrocytes, CLPs, naïve T cells, CD8^+^ T cells, Pro-B cells, B cells, dendritic cells, and NK cells (Fig. 2A). These cell types were characterized by markers such as *CD34*, *AVP*, *SPINK2*, *CD3E*, *HBA1*, *MPO*, *CD14*, *LYZ* and others (Fig. S2A). Furthermore, an increased occurrence of CMPs, GMPs, and monocytes was observed in diagnosis and relapse samples, while neutrophils were predominantly found in MRD samples (Fig. 2B, 2C). It was consistent with the clinical features. By comparing these AML hematopoietic cells with healthy bone marrow cells, we classified the AML hematopoietic cells into three types: cells with a cell state in AML (Cell^AML^), cells with an intermediate cell state (Cell^inter^) and cells with a cell state in healthy bone marrow (Cell^BM^) (Fig. S2B, S2C). Notably, Cell^AML^ were prevalent with immature cells, whereas Cell^BM^ were dominant with mature cells (Fig. S2D, S2E). As expected, we found Cell^AML^ were prevalent in diagnostic and relapse samples, while Cell^BM^ were dominant in MRD samples (Fig. S2E).

**Figure 2.**
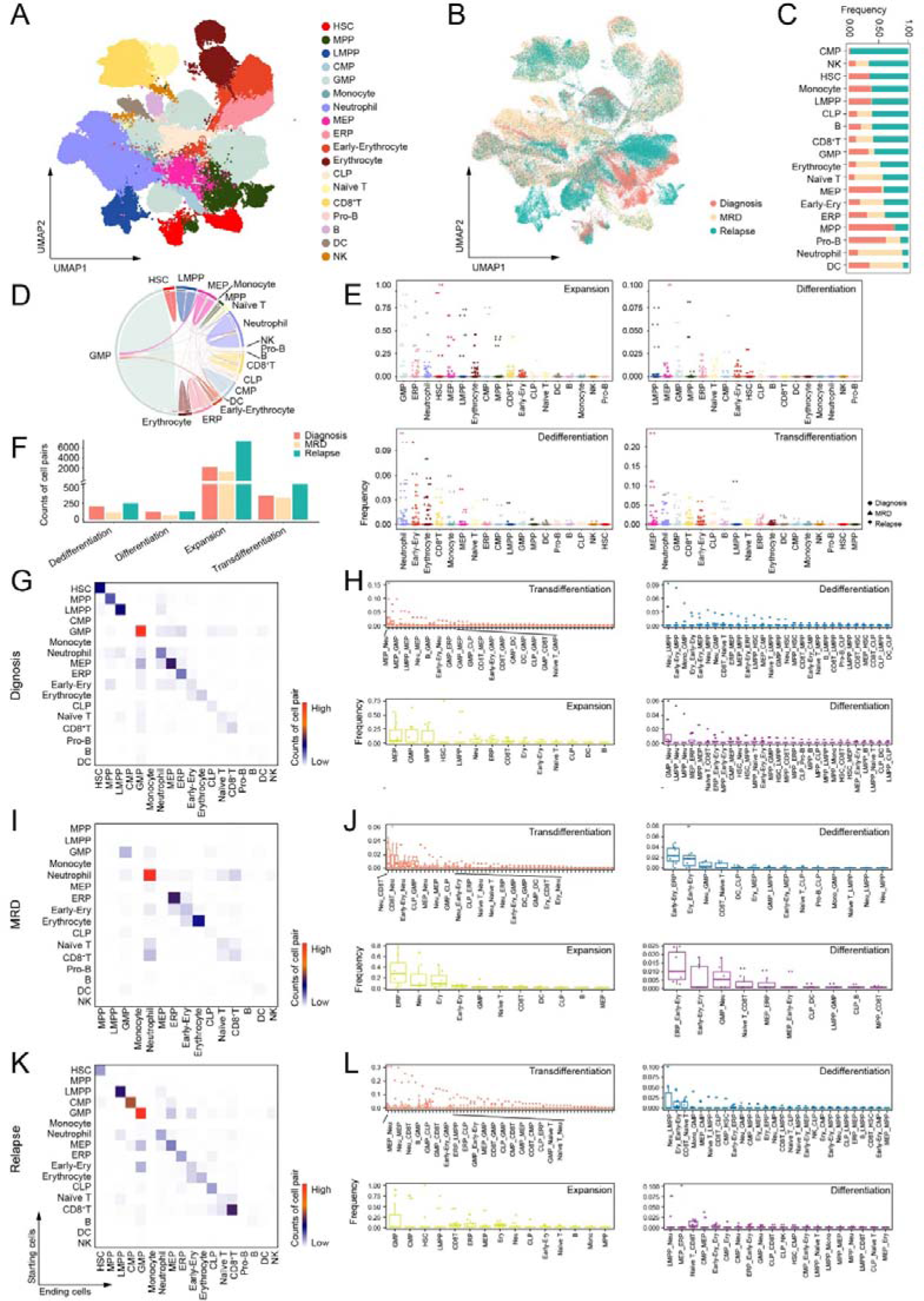
Characterization and dynamics of hematopoietic cells along with clinical process in AML. (A) UMAP plot showing the major lineages of hematopoietic cells in AML samples. (B) UMAP plot showing the clinical stage of AML hematopoietic cells. (C) Bar plot showing the clinical stage across the different cell types. (D) Circus plot demonstrating the dynamics of hematopoietic cells in the AML datasets. (E) Boxplots illustrating the frequency of cell changing types across the different starting cell types in (D). (F) Bar plot showing the counts of cell changing types across different clinical stages. (G) Heatmap showing the counts of cell changing types in the AML diagnosis samples. The cell types in the rows were the starting cells, and the cell types in the columns were the ending cells. (H) Boxplots illustrating the frequency of cell pairs across the different cell changing types in AML diagnosis samples. (I) Heatmap showing the counts of cell changing types in the AML MRD samples. (J) Boxplots illustrating the frequency of cell pairs across the different cell changing types in AML MRD samples. (K) Heatmap showing the counts of cell changing types in the AML relapse samples. (L) Boxplots illustrating the frequency of cell pairs across the different cell changing types in AML relapse samples.

According to the constructed lineage tracing atlas shown in circus plot (Fig. 2D, Table S2), a higher frequency of expansion and differentiation was observed in stem cells and progenitor cells, besides with the expansion of neutrophils and erythrocytes (Fig. S2F). In particular, it was shown that neutrophils, early-erythrocytes and erythrocytes as the top 3 could undergo dedifferentiation into immature cells, and LMPPs and early-erythrocytes were more likely to be generated by dedifferentiation from other cell types. MEPs, neutrophils and GMPs showed a higher propensity of transdifferentiation compared to other cells (Fig. 2E, Fig. S2G). Moreover, many differences in cell changings during the clinical process were identified. We found that counts of expansion dominated in all three clinical process samples, and counts of cell changings in MRD samples were fewer than that in diagnosis samples and relapse samples (Fig. 2F). In diagnosis samples, a high level of expansion in GMPs, LMPPs and MPPs and differentiation from these cells to neutrophils were observed. Transdifferentiation from MEPs to neutrophils and from MEPs to GMPs occurred frequently (Fig. 2G, 2H). Unlike in diagnosis and relapse samples, expansion occurred mainly in ERPs, neutrophils and erythrocytes, and differentiation occurred mainly from ERPs to erythrocytes, from early-erythrocytes to erythrocytes and from GMPs to neutrophils in MRD. Transdifferentiation from neutrophils to CD8^+^ T cells and vice versa were the top 2 in MRD (Fig. 2I, 2J). Of note, the frequency of transdifferentiation and dedifferentiation in MRD was far lower than that in diagnosis and relapse samples. Cell changings in relapse were similar to that in diagnosis. Apart from the high frequency of expansion in GMPs, there were transdifferentiation from MEPs to neutrophils and the opposite, and transdifferentiation from neutrophils to CD8^+^ T cells. The frequency of dedifferentiation from neutrophils to LMPPs was extremely high in relapse as well as in diagnosis samples (Fig. 2K, 2L). Taken altogether, these data suggested that the hematopoietic cells under AML bone marrow microenvironment acquired the plasticity to undergo cell fate changings and varied along with clinical process.

We found that the counts of cell changings between Cell^AML^ were predominantly in relapse and diagnosis samples, while total counts of cell changings in MRD samples were the least (Fig. S2H). We observed a notable preference in expansion of immature cells and transdifferentiation between MEPs and GMPs for cell changings among the Cell^AML^. The most cell changings from Cell^AML^ to Cell^BM^ and vice versa were both expansion of GMPs. The predominant cell changings between Cell^BM^ were mainly occurred in CD8^+^ T cells and Naïve T cells and erythroid cells, respectively (Fig. S2I). Additionally, the most cell changings between Cell^AML^ were expansion which probably resulted from the expansion of GMPs and the most cell changings from Cell^BM^ to Cell^AML^ were transdifferentiation. We observed a higher frequency of cell changings between Cell^AML^ were mostly in immature cells, as well as in diagnosis and relapse samples (Fig. S2J). These observations suggested that the dynamics of AML hematopoietic cells correlated with the clinical progress of AML.

### Dynamics of Hematopoietic Cells within CMML

We compiled about 40 CMML samples into a comprehensive dataset, where we discerned major hematopoietic cell types, including HSCs, MPPs, LMPPs, GMPs, MDPs, MKPs, CLPs, T cells, B cells, erythrocytes, monocytes and neutrophils using specific markers (Fig. 3A, Fig. S3A). Utilizing healthy bone marrow samples as a reference, we classified cells in CMML into three types containing cells with a cell state in CMML (Cell^CMML^), Cell^inter^ and Cell^BM^ based on the proportion of healthy bone marrow cells (Fig. S3B). The immature cells were mostly Cell^CMML^, similar to AML. However, the proportion of Cell^inter^ in CMML was higher than that in AML (Fig. S3C).

**Figure 3.**
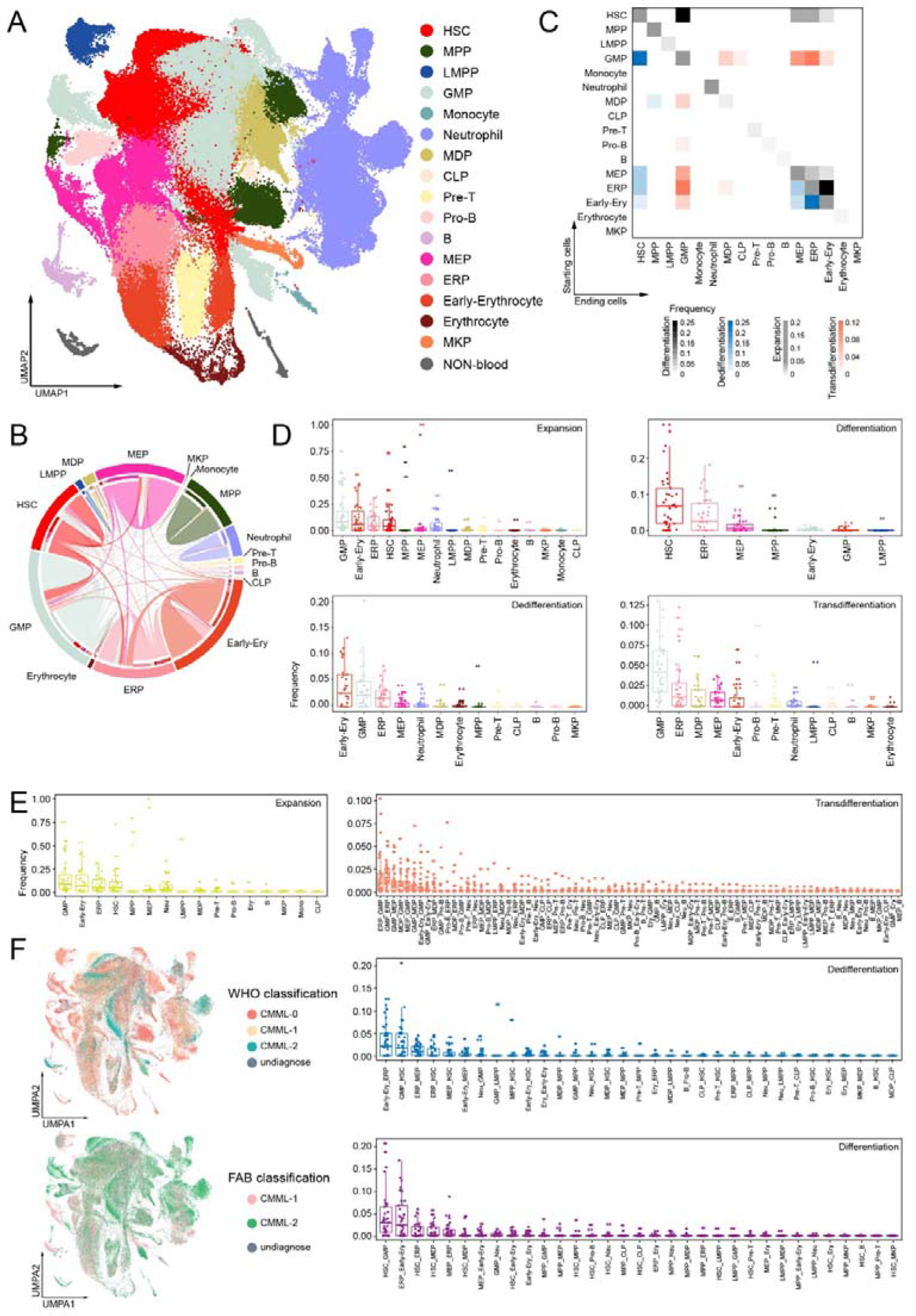
Characterization and dynamics of hematopoietic cells in CMML. (A) UMAP plot showing the major lineages of hematopoietic cells in CMML samples. (B) Circus plot demonstrating the dynamics of hematopoietic cells in the CMML datasets. (C) Heatmap showing the frequency of cell changing types in the CMML datasets analysis. The cell types in the rows were the starting cells, and the cell types in the columns were the ending cells. Different colors represented different cell changing types. The frequency was calculated through dividing the counts of each cell change type by the total number of all cell changing types. (D) Boxplots illustrating the frequency of cell changing types across the different starting cell types in (B). (E) Boxplots illustrating the frequency of cell pairs across the different cell changing types in AML samples. (F) UMAP plots showing the distribution of WHO classification and FAB classification.

Utilizing the previously established lineage tracing method, we observed that GMPs exhibited the higher dynamics (Fig. 3B, 3C, Table S3). GMPs showed the highest frequency of expansion and transdifferentiation as both the starting cells and the ending cells (Fig. 3D, Fig. S3D). Also, GMPs had a higher frequency of dedifferentiation as starting cells and differentiation as ending cells. Except for GMPs, ERPs and early-erythrocytes had a higher frequency of expansion (Fig. 3E). Also, high differentiation included cell changings from HSCs to GMPs and from ERPs to early-erythrocytes. The top 2 dedifferentiation were cell changings from early-erythrocytes to ERPs and cell changings from GMPs to HSCs and the top 2 trandifferentiation were cell changings between GMPs and ERPs. In conclusion, these data suggested that GMPs and erythroid cells under CMML bone marrow microenvironment acquire the plasticity to undergo cell fate changings.

Cell changings between Cell^CMML^ obtained high transdifferetiation and dedifferentiation among immature cells and erythroid cells besides expansion, while cell changings between Cell^BM^ mainly occurred in expansion (Fig. S3E). In total, cell changings between Cell^CMML^ had a higher frequency of differentiation and dedifferentiation, while cell changings from Cell^BM^ to Cell^CMML^ occurred mainly in dedifferentiation (Fig. S3F). In addition, the counts of cell changings between Cell^CMML^ were much higher than other cell changings, which also mainly occurred in immature cells. On the contrary, cell changings between Cell^BM^ happened mainly in mature cells. Combining with WHO classification and FAB classification (Fig. 3F), we found MEPs had more expansion in WHO0 and MPPs and HSCs had more expansion in WHO1 and WHO2. The phenomenon that the more immature cells were in the higher level of WHO subtypes was consistent with WHO classification (*29*) (Fig. S3G). We found that immature cells had more cell changings in FAB2 than in FAB1. The phenomenon that the more immature cells were in FAB2 than in FAB1 was consistent with FAB classification (*30*) (Fig. S3H).

### Less Dynamics of pDC Cells within BPDCN Microenvironment

BPDCN is a leukemia characterized by an elevated quantity of malignant pDCs (*31*). The UMAP revealed a significant presence of pDCs, as well as HSPCs, GMPs, monocytes, neutrophils, early-erythrocytes, erythrocytes, MKPs, B cells, T cells, and NK cells (Fig. 4A, Fig. S4A). Remarkably, there was a notable surge in NKT cells in relapse samples, suggesting a potential immune system response to the increasing pDC cell population (Fig. 4B, 4C) (*32*). CD8^+^ T cells and erythrocytes dominated in diagnosis samples. Based on healthy bone marrow hematopoietic cells, the majority of NK cells, NKT cells and pDC cells were identified as cells with a cell state in BPDCN (Cell^BPDCN^), while most T cells were Cell^inter^ (Fig. S4B, S4C). Other cell types were mainly categorized as Cell^BM^.

**Figure 4.**
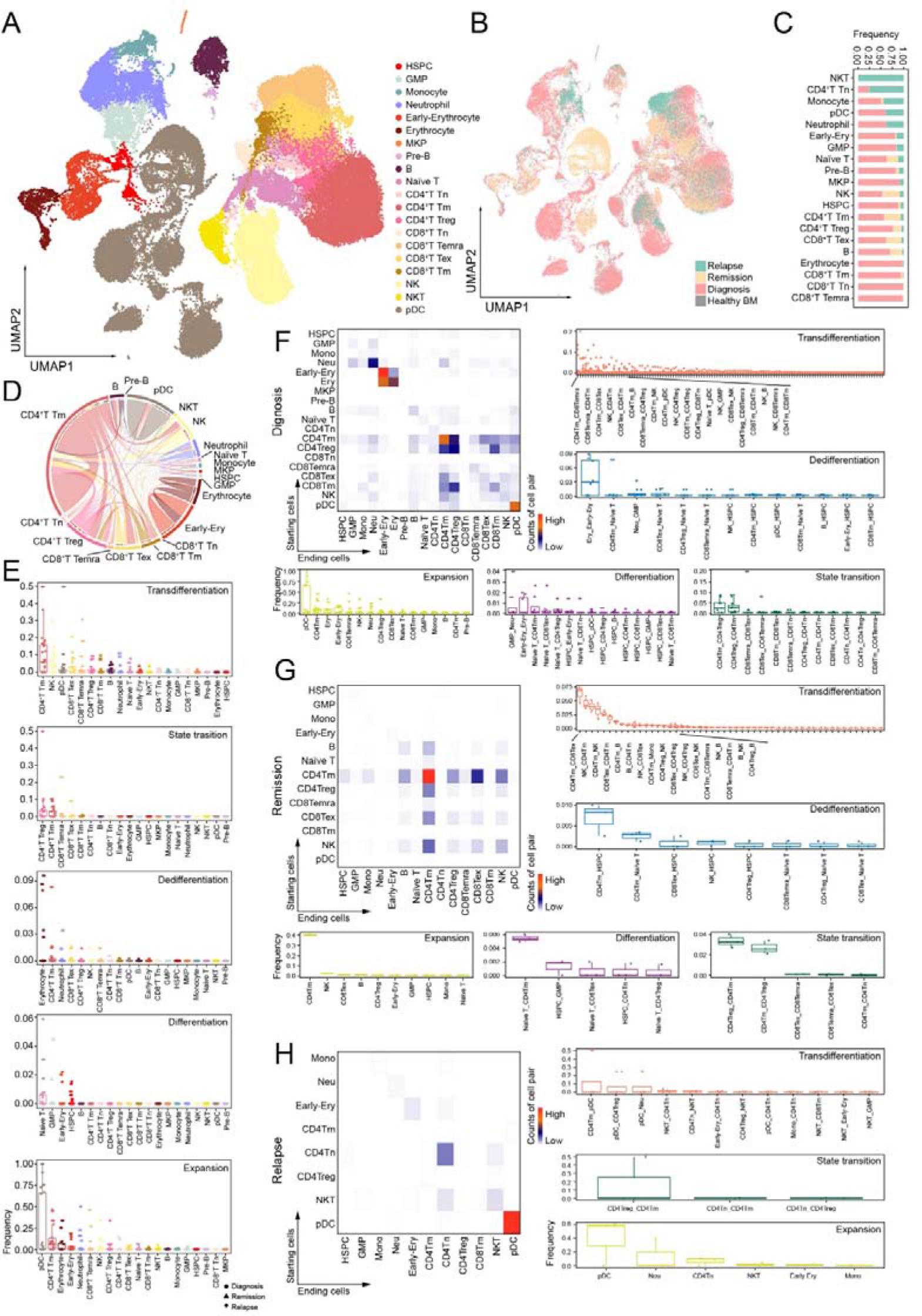
Less dynamics of pDC and high dynamics of CD4^+^T cells in BPDCN. (A) UMAP plot showing the major lineages of hematopoietic cells in BPDCN samples. (B) UMAP plot showing the integration of BPDCN and healthy bone marrow scRNA-seq datasets. (C) Bar plot showing the clinical stage across the different cell types. (D) Circus plot demonstrating the dynamics of hematopoietic cells in the BPDCN datasets. (E) Boxplots illustrating the frequency of cell changing types across the different starting cell types in (D). (F) Heatmap showing the counts of cell changing types in the BPDCN diagnosis samples. The cell types in the rows were the starting cells, and the cell types in the columns were the ending cells. Boxplots illustrating the frequency of cell pairs across the different cell changing types in BPDCN diagnosis samples. (G) Heatmap showing the counts of cell changing types in the BPDCN remission samples. Boxplots illustrating the frequency of cell pairs across the different cell changing types in BPDCN remission samples. (H) Heatmap showing the counts of cell changing types in the BPDCN relapse samples. Boxplots illustrating the frequency of cell pairs across the different cell changing types in BPDCN relapse samples.

Interestingly, pDCs exhibited minimal transdifferentiation from/to other hematopoietic cell types, with limited transdifferentiation from CD4^+^ T_m_ and CD4^+^ T_reg_ (Fig. 4D, Fig. S4D, Table S4), consistent with our previous findings in solid tumor immune microenvironments (*28*). The predominant cell changings observed in pDCs were expansions (Fig. 4E). Noteworthy is the plastic role of CD4^+^ T cells, especially CD4^+^ T_m_ and CD4^+^ T_reg_, in the BPDCN bone marrow microenvironment, displaying considerable transdifferentiation and state transition (Fig. 4E, Fig. S4E) (*33*). Along with clinical process, we found that counts of expansion were extremely high across different clinical samples and counts of transdifferentiatiton were high in diagnosis and remission samples (Fig. S4F). In diagnosis samples, the frequency of expansion in pDC was far higher than other expansion. The frequency of differentiation from early-erythrocytes to erythrocytes and dedifferentiation from erythrocytes to early-erythrocytes were also high. Additionally, T cells were active in cell changings, including transdifferentiation from CD4^+^ T_m_ to CD8^+^ T_emra_ and vice versa, transdifferentiation from CD4^+^ T_m_ to CD8^+^ T_ex_ and state transition from CD4^+^ T_m_ to CD4^+^ T_reg_ and vice versa (Fig. 4F). In remission samples, T cells, especially CD4^+^ T_m_, underwent dedifferentiation to HSPCs and state transition from/to CD4^+^ T_reg_. In addition, CD4^+^ T_m_ underwent transdifferentiation to CD8^+^ T_ex_ and from/to NK cells (Fig. 4G). In relapse samples, there was high frequency of expansion in pDC and neutrophils, state transition from CD4^+^ T_reg_ to CD4^+^ T_m_ and transdifferentiation from CD4^+^ T_m_ to pDC, as well as from pDC to CD4^+^ T_reg_ (Fig. 4H). In total, we discovered that the frequency of transdifferentiation and state transition were high in diagnosis and relapse samples and low in remission samples. Collectively, these findings indicated that T cells and NK cells displayed elevated dynamics, whereas pDC exhibited lower dynamics in BPDCN.

Within the BPDCN microenvironment, the predominant cell changings between Cell^BPDCN^ were expansion of pDC. There were minimal cell changes between Cell^BPDCN^ and Cell^BM^. Cell changings between Cell^BM^ occurred mainly in erythroid cells (Fig. S4G). As anticipated, cell changings between Cell^BPDCN^ exhibited in relapse samples and diagnosis samples, whereas cell changings between Cell^BM^ mainly exhibited in diagnosis samples (Fig. S4H). Apart from pDC cells, CD8^+^ T_ex_ and CD4^+^ T_reg_ were more prone to cell changings between Cell^BPDCN^, while CD4^+^ naïve T cells predominantly underwent cell changings from Cell^BM^ to Cell^BPDCN^. In addition, erythroid cells and myeloid cells underwent cell changings between Cell^BM^. Notably, cell changings of differentiation and dedifferentiation were both mainly between Cell^BM^, while cell changings of state transition were all between Cell^BPDCN^. Cell changings from Cell^BM^ to Cell^BPDCN^ were mainly occurred in transdifferentiation. (Fig. S4H). These observations indicated that the dynamics of BPDCN were primarily associated with pDCs ,T cells and NK cells.

### Different Dynamics of B Lymphoid Blast Cells within B-ALL and B-CLL Microenvironment

Distinct B lymphoid blast cells in both B-ALL and B-CLL were successfully identified, alongside B cells, T cells, erythroid cells and myeloid cells (Fig. 5A). Based on different sources of B lymphoid blast cells (Fig. 5B, 5C) and the specific markers *CD19*, *CD79A*, *DNTT*, *IGLL1* for B cells (Fig. S5A), these B lymphoid blast cells were further categorized into B-ALL blast cells, B-CLL blast cells and B-common blast cells which originated from both B-ALL and B-CLL. Using the healthy bone marrow data, we identified these hematopoietic cells into three cell types as cells with a cell state in lymphoblastic leukemia (Cell^Lym^), Cell^inter^ and Cell^BM^ (Fig. S5B). As anticipated, B lymphoid blast cells were predominantly identified as Cell^Lym^ (Fig. S5C). Our data revealed that CLL1 and CLL2 samples had a higher proportion of Cell^Lym^ (*34*), while the ALL remission samples (ALL1204 and ALL rem1) had a lower percentage of Cell^Lym^ (*35*).

**Figure 5.**
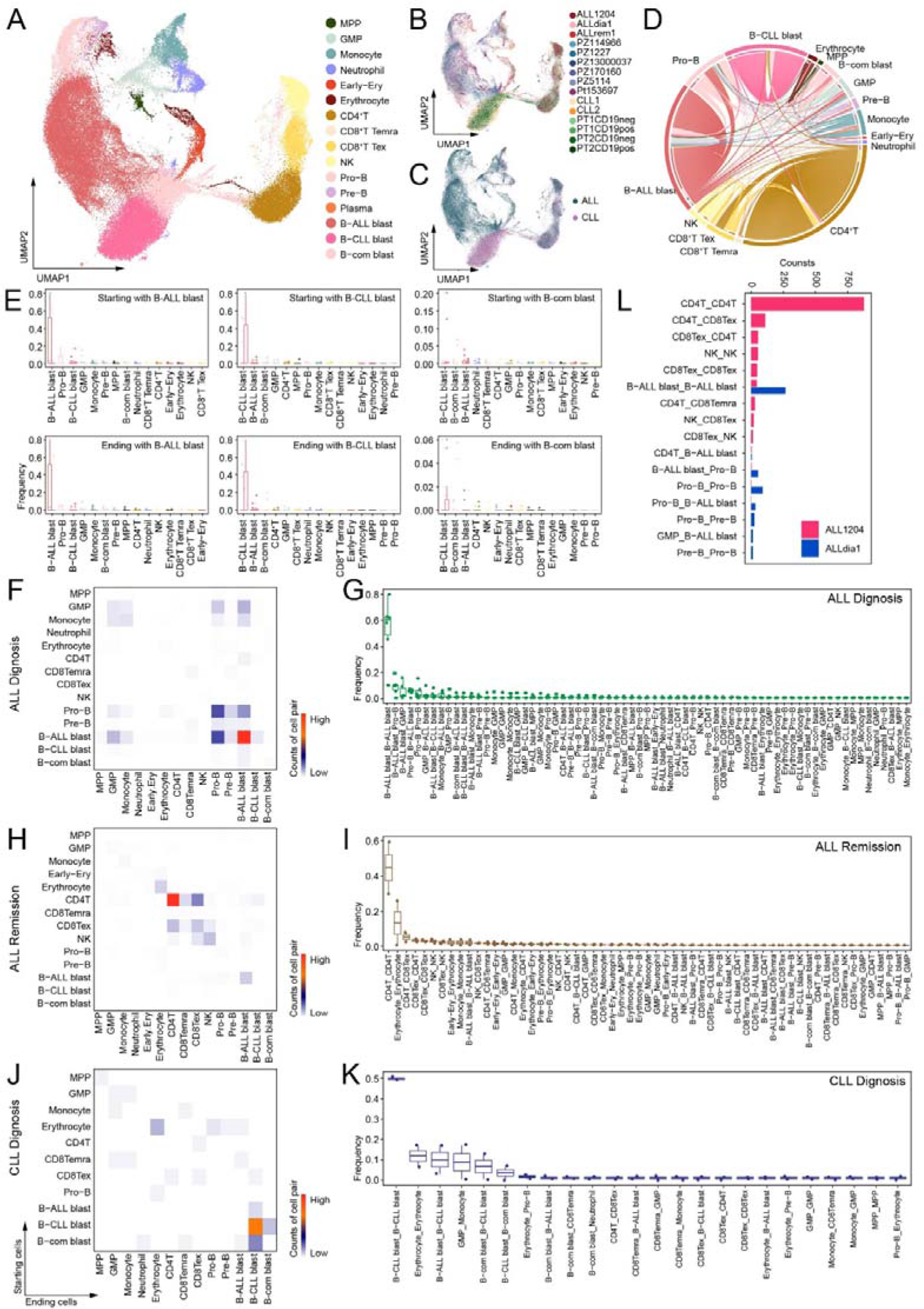
Distinct dynamics of hematopoietic cells in the integration of ALL and CLL datasets. (A) UMAP plot showing the major lineages of hematopoietic cells in lymphoblastic leukemia samples. (B) UMAP plot showing the distribution of lymphoblastic leukemia samples. (C) UMAP plot showing the distribution of ALL and CLL. (D) Circus plot demonstrating the dynamics of hematopoietic cells in the lymphoblastic leukemia datasets. (E) Boxplots illustrating the frequency of the cell changing starting with or ending with the three B leukemia blast cell types. (F) Heatmap showing the counts of cell changing types in the ALL diagnosis samples. The cell types in the rows were the starting cells, and the cell types in the columns were the ending cells. (G) Boxplots illustrating the frequency of cell pairs in ALL diagnosis samples. (H) Heatmap showing the counts of cell changing types in the ALL remission samples. (I) Boxplots illustrating the frequency of cell pairs in ALL remission samples. (J) Heatmap showing the counts of cell changing types in the CLL diganosis samples. (K) Boxplots illustrating the frequency of cell pairs in CLL diganosis samples. (L) Bar plot displaying the counts of cell pairs in ALLdia1 and ALL1204 samples.

B-ALL blast cells and B-CLL blast cells exhibited a higher frequency of expansions compared to B-common blast cells (Fig. 5D, Fig. S5D, Table S5). Notably, B-ALL blast cells were the primary cell type to which other cell types transdifferentiated. B-common blast cells showed a tendency to transition to both B-ALL blast cells and B-CLL blast cells, while B-ALL blast cells and B-CLL blast cells exhibited a reciprocal relationship with B-common blast cells (Fig. 5E). This may explain the transformation between ALL and CLL disease (*36*). These transformation demonstrated the plasticity of B cells in B lymphoid leukemia (*37*), which was not observed in other types of leukemia mentioned earlier. In ALL diagnosis samples containing our data ALLdia1, cell changings mainly occurred among Pro-Bs, Pre-Bs and B-ALL blast cells (Fig. 5F). Expansion of B-ALL blast cells, transdifferentiation from B-ALL blast cells to Pro-Bs and to GMPs were the top 3 high frequency of cell changings (Fig. 5G). In ALL remission samples including our data ALLrem1 and ALL1204, T cells played an important role in immune function, containing expansion of CD4^+^ T and transdifferentation from CD4^+^T to CD8^+^ T_ex_ (Fig. 5H, 5I). In CLL diagnosis samples of our data CLL1 and CLL2, cell changings mainly involved the expansion of B-CLL blast cells and erythrocytes, differentiation from GMPs to monocytes and transformation from B-ALL blast cells to B-CLL blast cells and from B-common blast cells to B-CLL blast cells (Fig. 5J, 5K). As our data ALLdia1 and ALL1204 were diagnosis and remission samples of one patient, we compared the cell changings of these two clinical stages (Fig. 5L). Expansion of B-ALL blast and CD4^+^ T were the most frequent cell changings in ALLdia1 and in ALL1204 respectively, which were consistent with clinical process. Overall, these data illustrated the distinct dynamics and plasticity of B lymphoid blast cells in ALL and CLL within bone marrow microenvironment.

Cell changings between Cell^Lym^ mostly involved in B lymphoid blast cells, Pro-Bs and Pre-Bs, while cell changings between Cell^BM^ mostly involved T cells, NK cells, erythrocytes, monocytes and GMPs (Fig. S5E, S5F). We found that cell changings from Cell^Lym^ to Cell^BM^ were predominantly transformation from B lymphoid blast cells to other cell types. On the contrary, cell changings from Cell^BM^ to Cell^Lym^ dominated in transformation from other cell types to B lymphoid blast cells. In addition, ALL diagnosis samples had more cell changings between Cell^Lym^, while CLL diagnosis and ALL remission samples obtained more cell changings between Cell^BM^. B-ALL blast cells, B-CLL blast cells and B-common blast cells were the top 3 cell types undergoing cell changings between Cell^Lym^ (Fig. S5F). As a whole, these observations showed that the dynamics and plasticity of ALL and CLL were mainly associated with B lymphoid blast cells.

### Potential Mediators in Bone Marrow Microenvironment Contributing to Hematopoietic Cell Plasticity

To investigate the extrinsic factors influencing cell changings, we conducted a comparison between cells undergoing expansions and those undergoing differentiation, state transition, transdifferentiation and dedifferentiation, respectively, using Nichenet (*38*). Ligand-receptor analysis revealed selective interactions of these cells with the bone marrow hematopoietic microenvironmental cells, including cells with a cell state in leukemia (Cell^Leu^) and healthy bone marrow hematopoietic cells. These interactions were influenced by soluble factors secreted by these cells (Fig. 6A, Table S6). The ligands TGFβ1, ICAM1, TNF and IL1β were identified in different cell changing types across various leukemias. It had been reported that TGFβ1 has the ability to drive cell reprogramming (*39, 40*), which may play a potential role in cell changings within leukemia microenvironment. The ligands ANXA1, IL1β, LGALS3 and HMGB1 were mainly identified in AML and CML. ANXA1 may promote the tumor progression through inhibiting neutrophil tissue accumulation and activating neutrophil apoptosis so that it may induce hematopoietic cells reprogramming (*41*), as well as HMGB1 (*42*). In addition, we made a comparison between the cells undergoing expansion and other changings (Fig. S6A). The ligands TNFSF13β, ITGA4, CLEC11A and PTDSS1 were identified mainly in cell changings of immature erythroid cells within AML and CML microenvironment, as well as cell changings of Pro-Bs in ALL. The ligands IFNγ and IL1α worked on GMPs in AML and the ligands LTB, TNFSF10, IL7 and TNF worked on myeloid cells in AML and CML microenvironment. The ligands TNFSF12, HGF and PIK3CB were mainly identified in cell changings of the stem and progenitor cells. The ligands DUSP18, ITGB2 and GSTP1 were mainly identified in cell changings of lymphoid cells. Then we computed the number of the potential ligands that were either specific to a cell type or shared among multiple cell types (Fig. 6B). CD4^+^ T_m_ had the most specific ligands and these ligands drove the state transition in BPDCN, while ERP and GMP had both 3 specific ligands. Neutrophil, GMP and MEP had top 3 quantities of total potential ligands. According the ligand-receptor analysis, we found that in AML and CML, immature cells had more tendency to undergo cell changing. In BPDCN, cells had less dynamics within microenvironment, while in ALL and CLL, lymphoid cells made more cell changings (Fig. 6C). This is consistent with our previous conclusion. Further inspection of extrinsic factors revealed that CCL5 was significant in non-expansion changing and in Cell^Leu^, while CCL17 was significant in expansion changing and in Cell^Leu^. Additionally, GSTP1 was significant in Cell^Leu^ (Fig. S6B). CCL5 had been implicated in leukemia progression and tumor malignancy. Inhibition of CCL5 with maraviroc had been shown to alleviate leukemia progression, and survival curves indicated a decrease in survival probability associated with CCL5 and GSTP1 (Fig. S6C) (*43–45*). Similarly, the survival curves of CCL17 and GZMB demonstrated a negative influence of these ligands on leukemia. In conclusion, we identified several extrinsic factors that could potentially influence the dynamics of leukemia hematopoietic cells within the microenvironment.

**Figure 6.**
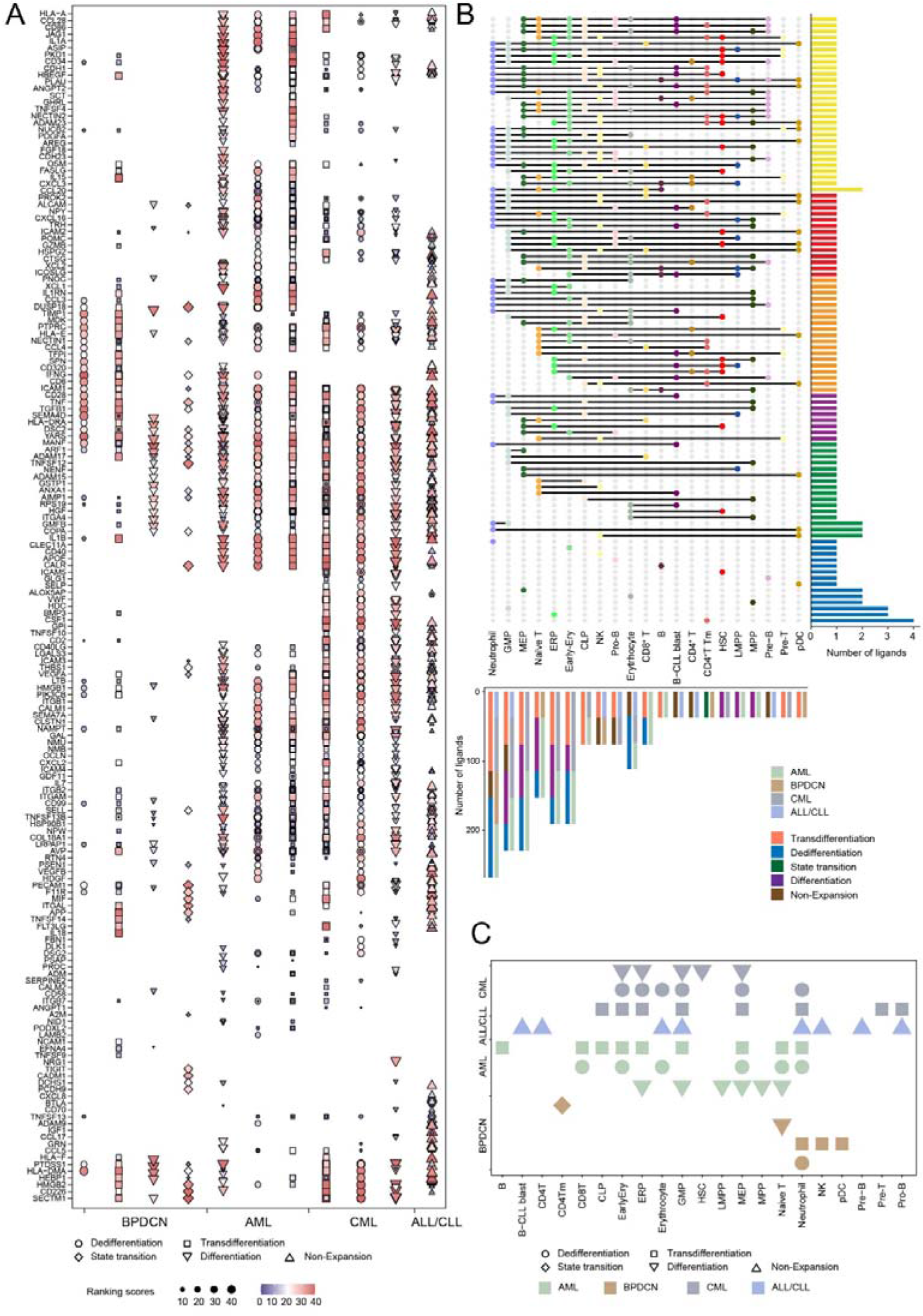
Factors correlated with the leukemia bone marrow microenvironment. (A) Dot plot illustrating the potential ligands driving the different cell changing types across various leukemias within the bone marrow microenvironment. Different patterns represented different changing types. (B) Number of the potential ligands in (A) following each cell type, grouped by sharing pattern, either specifically driving one cell type or shared by two or more cell types (top). Bar plot showing the number of these ligands across various leukemia types and cell changing types, respectively (bottom). (C) Heatmap displaying the plasticity of different cell types in various leukemia types. Different patterns represented different changing types.

### Specific Transcriptional Regulations Controlling the Reprogramming of Hematopoietic Cells

SCENIC analysis, which reconstructs regulons (TFs and their target genes), was employed to infer the regulatory networks of hematopoietic cells (*46*). Based on the SCENIC AUC score, it was observed that Cell^Leu^, Cell^inter^ and Cell^BM^ were clustered, respectively (Fig. S7A). To identify the regulons influencing cell changings, a comparison of regulons in changing cells across leukemia samples was conducted (Fig. 7A). In AML, CMML, and BPDCN, regulons, including KLF3(+), E2F2(+), MXI(+), GATA1(+), KLF1(+) and NFE2(+), positively correlated with dedifferentiation. The transcriptional factor KLF3 and E2F2 had been proved that they play an important role in cell reprogramming (*47, 48*). Notably, these regulons exhibited minimal expression in healthy bone marrow samples and higher expression in CMML, ALL, and BPDCN (Fig. 7B). The regulons SOX4(+), ERG(+), MYB(+) and ETS2(+) were positively correlated with all cell changings from B-ALL blast cells to Cell^Lym^, Cell^inter^ and Cell^BM^, while CEBPE(+) and FOS(+) were higher in cell changings from B-ALL blast cells to Cell^BM^ (Fig. 7C). The regulons EOMES(+), RELB(+) and RUNX3(+) were negatively correlated with cell changings from B-ALL blast cells to Cell^inter^ and Cell^Lym^ and the regulons ETS1(+) and PRDM1(+) were negatively correlated with cell changings from B-ALL blast cells to Cell^BM^ and Cell^Lym^. However, the regulons RELB(+), POU2F2(+), IRF8(+) and SPIB(+) were positively correlated with all cell changings from B-CLL blast cells to Cell^Lym^, Cell^inter^ and Cell^BM^, while the regulons ETS2(+) and MYB(+) were negatively correlated with cell changings from B-CLL blast cells to Cell^Lym^, Cell^inter^ and Cell^BM^. The regulons, such as ETS2(+), MYB(+) and RELB(+), worked oppositely on the cell changings in ALL and CLL.

**Figure 7.**
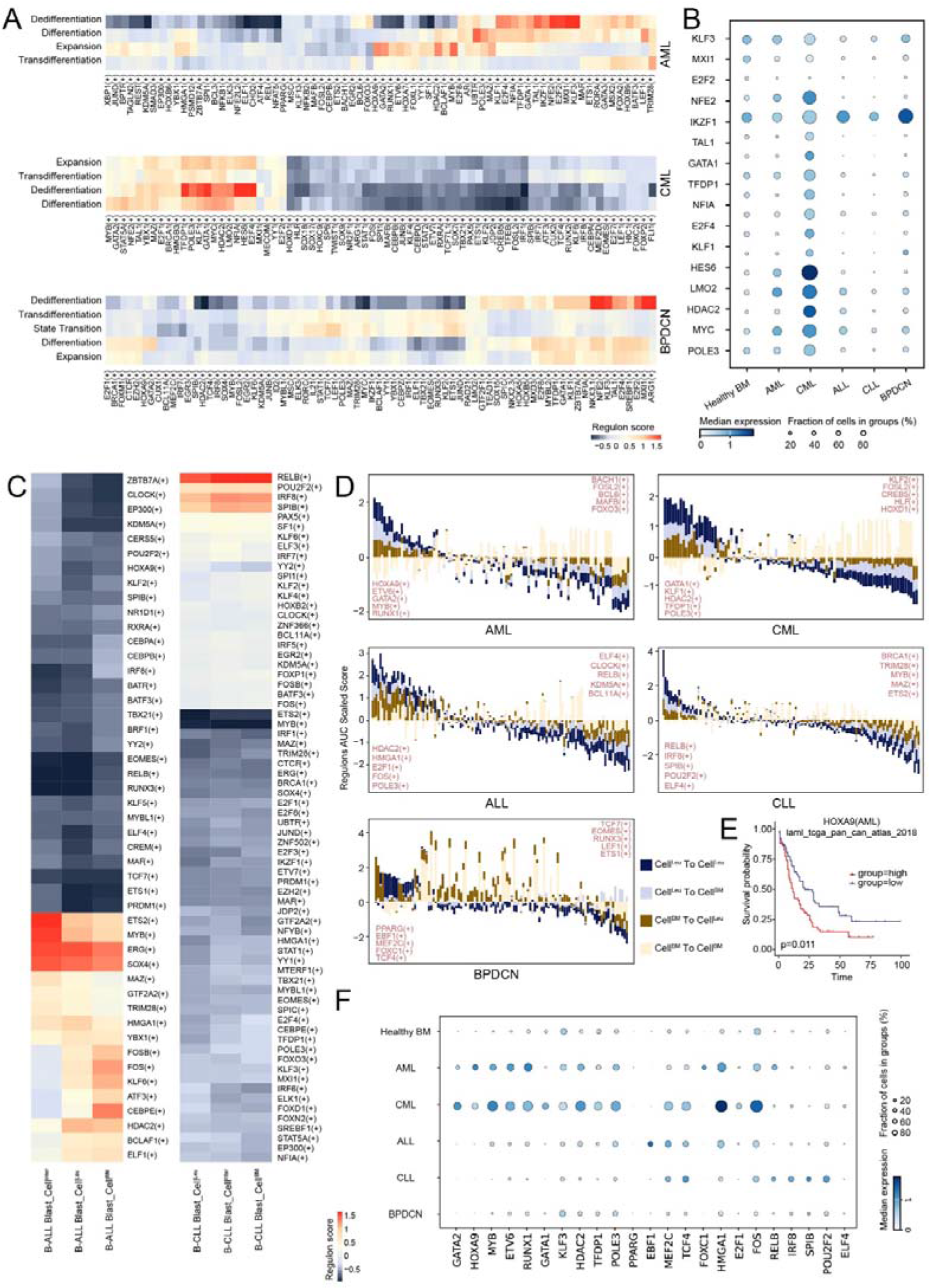
Regulons affecting the cell changings across various leukemias. (A) Heatmap showing the regulons across the different cell changings in AML, CML and BPDCN datasets. (B) Dot plot illustrating the expression of the partial transcriptional factors in (A) across various leukemias and healthy bone marrow. (C) Heatmap showing the regulons of the cell changings from B leukemia blast cells to Cell^Leu^ (left) and from Cell^Leu^ to B leukemia blast cells (right). (D) Bar plots displaying the regulons AUC scaled score in the cell changings among Cell^Leu^ and Cell^BM^ across the different leukemia types, which were ordered by cell changings from Cell^Leu^ to Cell^Leu^ and cell changings from Cell^BM^ to Cell^Leu^ . The top regulons and the bottom regulons were displayed on the left and right, respectively. (E) Kaplan-Meier plots showing worse clinical outcome in AML patients with the higher expression of HOXA9. (F) Dot plot illustrating the expression of the top 5 transcriptional factors in (D) across various leukemias and healthy bone marrow.

Then we compared the regulons of the cells that changed into myeloid cells, erythroid cells and lymphoid cells, respectively. Compared with the myeloid cells undergoing expansion, we found that both CD8^+^ T_emra_ cells changed into GMPs in CLL and NK cells changed into monocytes in CLL shared similar regulons, which implied that these two cells may be undergoing the intermediate cell state in the cell changings into myeloid cells (Fig. S7B). Compared with the erythroid cells undergoing expansion, we found Pre-Bs, Pro-Bs, HSPCs and neutrophils, which changed into erythrocytes, early-erythrocytes and ERPs in ALL, CLL, BPDCN and CML, respectively, shared similar regulons. This implied that these cells may be maintaining the intermediate cell state in the cell changings into erythroid cells (Fig. S7C). Unfortunately, compared with the lymphoid cells undergoing expansion, we didn’t find the intermediate cell state in the cell changings to lymphoid cells (Fig. S7D). Taken together, the intermediate cell state observed in cell changings validated the dynamics of leukemia hematopoietic cells within the leukemia microenvironment.

The top five regulons and the bottom five regulons of cell changings between Cell^Leu^ and Cell^BM^ across leukemia samples were listed (Fig. 7D). The regulons were ordered by cell changings from Cell^Leu^ to Cell^Leu^ and from Cell^BM^ to Cell^Leu^. The survival curve of HOXA9, one of the top five regulons in AML, aligned with the conclusions drawn from the analysis (Fig. 7E). These top regulons did not express in healthy bone marrow samples which explained that these top regulons were positively with leukemia progression (Fig. 7F).

### PAGA implying the transformation between B-common blast cells and T cells

We conducted PAGA analysis on the transferring cells in all the above leukemia samples (Fig. S8A, S8B). Surprisingly, unlike B-ALL blast cells and B-CLL blast cells, we found that B-common blast cells exhibited higher connectivity scores with T cells and NK cells (Fig. S8C, Table S7). This led us to speculate that B leukemia cells might have the potential to transition into T leukemia cells (*49*). Notably, the correlation between B-common blast cells and T cells was higher than that between B-ALL blast cells and B-CLL blast cells (Fig. S8D). According to our previous findings, the frequency of transformation from B-common blast cells to CD8^+^ T cells was higher than other two cell types (Fig. S8E, S8F). This finding provides a valuable direction for our further investigations in leukemia research.

## Conclusion and Discussion

In this comprehensive study, we assembled a diverse collection of bone marrow hematopoietic cells spanning five leukemia types, delving into the intricate plasticity and dynamics of these cells within the leukemia bone marrow microenvironment. Leveraging lineage tracing based on SNP and mtDNA mutations inferred from scRNA-seq data, combined with accurate cell type definitions, we elucidated the cell changings of hematopoietic cells within multiple leukemia samples. Our results demonstrate that hematopoietic cells could acquire diverse plasticity and exist at different dynamic state across various leukemias. In AML, GMPs and neutrophils not only exhibited relatively higher dynamic potential to transition into multiple cell types, but also obtained a higher quantity of transitions from other cell types. Dynamics of hematopoietic cells in AML changed along with clinical process. The frequency of transdifferentiation in diagnosis and relapse samples was higher than that in MRD samples, while the frequency of expansion in diagnosis and relapse samples was lower than that in MRD samples. In CMML, besides GMPs existing in high dynamic states, ERPs and early-erythrocytes displayed a tendency to dedifferentiate into immature erythroid cells. Notably, the cell changings between Cell^AML^ or Cell^CMML^ primarily occurred in immature cells in both AML and CMML. In BPDCN, pDCs, mainly characterized as Cell^BPDCN^, seemed to maintain a more stable fate, showing minimal transitions to other cell types. In diagnosis samples, cell changings occurred mainly in lymphoid cells and pDC. In remission samples, only a few subtypes of T cells obtained higher dynamics and in relapse samples, there were extremely high quantity of expansion of pDCs. In B-ALL and B-CLL diagnosis samples, B-ALL blast cells and B-CLL blast cells emerged at the highest dynamic state, respectively. In ALL remission samples, CD4^+^T and erythrocytes showed high dynamics. In addition, we found B-common blast cells occurring both in ALL and CLL underwent transformation with B-ALL blast cells and B-CLL blast cells. The distinct behaviors of hematopoietic cells within various leukemia microenvironments illuminate the complex dynamics of these cells in leukemia contexts, providing valuable insights for further studies in the field.

Cell changings were highly influenced by extrinsic factors involved within complex leukemia bone marrow microenvironment, such as ANXA1, HMGB1, TNFSF13β, IL1β, TGFβ, TNF, and IFNγ. These factors, previously reported to impact cell fate to induce reprogramming, especially epithelial-mesenchymal-transition (*39, 40*) , played a crucial role in shaping the dynamics of cell changes. Each lineage cell types owned their specific ligands, for instance, TNFSF13β to erythroid cells, IFNγ to myeloid cells and GSTP1 to lymphoid cells. Notably, CCL5 emerged as significant in non-expansion changing and Cell^Leu^. Inhibition of CCL5 with maraviroc has been demonstrated to alleviate leukemia progression. Transcriptional factors were also identified to control cell changings. Regulons including KLF3(+), E2F2(+) and GATA1(+) were identified to influence the dedifferentiation and regulons including HOXA9(+), SOX4(+) and MYB(+) were identified to influence the transformation of B leukemia blast cells. Additionally, we found some cells undergoing myeloid cells and erythrocytes maintained intermediate state in cell changings, respectively. This may uncover the dynamics of hematopoietic cells during the process of reprogramming. Lastly, the connectivity score derived from PAGA suggested that B-common blast cells harbored the potential ability to transition into T leukemia blast cells. Additional exploration is necessary to clarify whether these influencing factors operate independently or synergistically.

Previous study had proved that hematopoietic cells within leukemia microenvironment could obtain plasticity to undergo reprogramming. Under conditions of hypoxia, B lymphoma cells are capable of converting to erythroblast-like cells (*17*). Lentiviral overexpression experiments demonstrated that ADAR1 promotes expression of the myeloid transcription factor PU.1 and induces malignant reprogramming of myeloid progenitors (*50*). Primary human BCR–ABL1+ B-ALL cells could be induced to reprogram into macrophage-like cells *in vitro* (*51*). Apart from these isolated cases, our study offered a comprehensive perspective on potential hematopoietic cell alterations in various types of leukemia. It not only validates previous findings but also unveils previously undiscovered changes.

The current gold standard for lineage tracing relies on genetic labelling through either fluorescent protein or genetic barcodes. Unfortunately, this method is limited to *in vitro* assays or animal models, making it challenging to utilize for human sample analysis (*52*). Single-cell RNA and DNA sequencing analysis of the same cell is considered to be the optimal technique for lineage tracing analysis. However, this method is often expensive and low-throughput, which prevents widespread use (*53*). To solve this problem, multiple bioinformatic methods concluding RNA velocity, Monocle pseudotime analysis and PAGA analysis were developed for lineage tracing (*54*). However, these methods are not precise enough for lineage tracing because they only focus on transcriptional information. And most of them didn’t consider cell reprogramming. SNP and mtDNA mutation inferred from scRNA-seq data individually have been proved to construct phylogenetic trees. Here, we combined them together as cell genetic barcodes and built strict filter criteria to uncover cell changings within leukemia bone marrow microenvironment. Nonetheless, future efforts to confirm other cell changings through canonical lineage tracing methods are still necessary.

In conclusion, our extensive analysis had advanced the current understanding of plasticity and the dynamics of leukemia bone marrow hematopoietic cells, providing a holistic perspective. This illumination extended to insights into the leukemia microenvironment, encompassing both local and systemic responses. We anticipated that the wealth of data generated in this large-scale study will contribute to a deeper understanding of the leukemia microenvironment and served as a catalyst for advancing clinical immunotherapy approaches in leukemia (*55*).

## Acknowledgements

We thank all the other stuffs in Shanghai Institute of Hematology and SJTU HPC for providing help on this project.

## Authors’ contributions

LC contributed to conception and design; CW contributed to single-cell sequence analysis and wrote the whole manuscript text and all figures and tables; YZ and SX contributed to experiment performance; All authors read and approved the final manuscript.

## Funding

This work is supported by the fund from Shanghai Municipal Health Commission (2022XD050), the fund from Shanghai Municipal Education Commission (22SG12) and the National Key R&D Program of China (2023YFC2508900).

## Data Availability

Sequence data that support the findings of this study have been deposited in the National Genomics Data Center with the primary accession code PRJCA022859.

## Competing Interest declaration

The authors declare that they have no competing interests.

## Materials and methods

### Single-cell RNA-Seq datasets in this study

We collected raw single-cell sequencing data on leukemia bone marrow hematopoietic cells from approximately 50 samples across 6 projects, which were diagnosed as one of the 5 leukemia types (Table S1). 5 leukemia samples were newly generated by our own including 3 ALL samples and 2 CLL samples. Our team obtained the data by downloading them from the GEO using SJTU HPC. We only selected data that met the following criteria: it had sequenced information on hematopoietic cells, and the raw sequence data or bam files were available for download. Healthy bone marrow data was also downloaded for the further verification and analysis.

### Sample collection

For scRNA-seq analysis, bone marrow (BM) samples were procured from a cohort comprising one patient diagnosed with acute common B cell lymphoid leukemia, two patients with chronic lymphoid leukemia, and one patient who had attained complete remission following treatment for acute lymphoid leukemia. The diagnoses of acute lymphoid leukemia and chronic lymphoid leukemia were established in accordance with The 5th edition of the World Health Organization classification of hematolymphoid tumors: Lymphoid Neoplasms. Specifically, patient sample ALLdia1 was obtained during the initial diagnosis of acute lymphoid leukemia, while patient sample ALL1204 was collected from the same individual after one cycle of induction therapy involving Vincristine sulfate, Idarubicin Hydrochloride, and dexamethasone (DVP). Patient sample ALLrem1 was acquired from a patient with acute lymphoid leukemia after seven cycles of chemotherapy, achieving complete remission as confirmed by both marrow smear examination and flow cytometry detection. Samples CLL1 and CLL2 were gathered from two patients at the time of initial chronic lymphoid leukemia diagnosis. The demographic and clinical characteristics of all patients are detailed in Table S1. These patients are all hospitalized between November 2023 and December 2023 in the Department of Hematology at Ren Ji Hospital.

### Single-cell RNA-seq data processing

All upstream analysis processes for scRNA-seq data were carried out on the SJTU HPC platform. Fastq files or bam files were obtained from the Siyuan No.1 platform using the wget function, SRA toolkit, or Aspera software. Bam files were then converted to the fastq file format using the Cell Ranger (version 3.1.0) bamtofastq function. To obtain bam files and expression matrixs, we processed the fastq files through the 10× scRNA-seq Cell Ranger Count pipeline using the GRCh38 human reference genome. Only quality-passed cells that passed the Cell Ranger filtering step were included in downstream analysis.

### Dimension Reduction and Unsupervised Clustering (Seurat)

The single-cell RNA expression matrixs underwent dimension reduction and unsupervised clustering using the Seurat pipeline (version 4.1.1) on RStudio at the HPC studio visualization platform. The cells were normalized and scaled for homogeneity, and highly variable genes were selected for downstream analysis. Principal component analysis (PCA) was performed on the scaled data to reveal the main axes of variation and denoise the data. Clustering of single cells by expression matrix was achieved by setting the parameter “resolution” of the FindClusters function to 2, generating more clusters for downstream cell type identification. Data were further reduced using the Uniform Manifold Approximation and Projection (UMAP) implemented in the RunUMAP function with default parameters. Cluster-specific marker genes were identified using the FindAllMarkers function with default parameters. To identify cell types, unsupervised clustering was performed on the hematopoietic cells to characterize the subsets of hematopoietic cells, identifying erythroid cells using *HBA1*, lymphoid cells using *CD3D* and myeloid cells using *CD14* following the above strategy. Subdivided cell types were defined by more specific markers published in the study.

### Dimension Reduction and Unsupervised Clustering (Scanpy)

Scanpy, an analysis toolkit suitable for massive single-cell sequence in python, was used for undergoing dimension reduction and unsupervised clustering and reconstructing the differentiation trajectory. The preprocessed feature-barcode matrix of leukemia hematopoietic cells was transferred into loom format to reconstruct the trajectory in Scanpy. The mean value of highly variable genes varied from 0.0125 to 3. PCA was computed based on the chosen variable genes. After preprocessing and PCA, the cells were clustered by Leiden algorithm. Differential marker genes were calculated for the further definition. The connectivity of the relationship of the cell types was constructed by partition-based graph abstraction (PAGA) algorithm (Table S6). The integration of data in scanpy was used with BBKNN, which could mix cells homogeneously.

### Integration of Multiple scRNA-seq Datasets by Harmony

We performed Harmony (version 0.1.1) analysis to integrate multiple scRNA-seq datasets and merge cells of the same type to eliminate batch effects caused by different projects. The output of the Harmony integration performed well in mixing the hematopoietic cells and was used as the input data for downstream analysis.

### Detection of Mitochondrial Mutations by Pysam

The bam files generated by Cell Ranger were imported into the Pysam (version 0.21.0) module using a custom Python script. The “pysam.AlignmentFile” function was used to transfer the bam file into the alignment file, allowing us to conveniently iterate over all of the read mappings in a specified region. We detected the mitochondrial information using the “pileup-engine” function.

To identify mitochondrial mutations, we aligned this sequence data with the human mitochondrial reference genome downloaded from the Ensemble website at the same mitochondrial position site. Allele frequency was calculated as the reads of a specific base at a position divided by the total reads at that position. The data of cells’ mitochondrial mutations were prepared for the next lineage tracing and further analysis.

### Detection of SNP by samtools and bcftools

To obtain SNP (Single Nucleotide Polymorphism) information for each cell, a series of steps were taken. First, the whole data bam was split into smaller bams for each cell using samtools. Then, samtools pileup was used to align the split bam files to the GRCh38 human reference genome. Next, bcftools (version 1.17) was employed to filter unreliable SNPs, which included those with a quality score of less than 10, a DP (read depth) score of less than 5, and SNPs near INDELs within 5 sites. The output files for each cell were merged into a single VCF file for further analysis to trace the lineage of cells in the tumor.

### Prediction of cell relationships through mitochondrial mutations

To ensure accurate lineage tracing, we filtered out variants with an allele frequency less than 5% to eliminate those that may disrupt heteroplasmy due to sequencing errors or RNA editing. We had tested various cutoff values and found 5% to be appropriate for our analysis. After filtering, we obtained new mitochondrial mutation data for each sample, and we analyzed the data in parallel and obtained results quickly.

We computed the correlation between every two cells by building a matrix that contains their mitochondrial mutations and frequencies. To obtain preliminary cell change results, we filtered correlation values less than 0.8, which we found to be a convincing cutoff to produce reasonable and beneficial lineage tracing results. We hypothesize that downstream cells should have more mitochondrial mutations than upstream cells since decedent cells inherit the ancestor’s mutation points and accumulate additional mutations. Therefore, we calculated the mitochondrial mutation number of cell pairs and defined the cell with more variants as the downstream cell and the cell with fewer mutations as the upstream cell.

### Prediction of Cell Relationships by SNP

The VCF files from each single cell were merged into a single file to obtain a precise result of cell relationships. To reduce interference, we filtered the SNPs a second time based on their quality score, removing those with a score less than 30. Additionally, we filtered out SNPs with insufficient read counts, with cutoff values ranging from about 10 to 600 depending on the sample. Next, we ran the same python script as in the previous step to obtain the correlation values between every two cells. We obtained preliminary cell change results calculated by SNP after filtering out pairs with a correlation value less than 0.6. We arrived at this cutoff value after extensive testing, as it provided reasonable and beneficial results for lineage tracing.

### Integration of the Two Relationships

To obtain a more accurate and precise result of cell change relationships, we merged the predictions obtained from mitochondrial mutation and SNP data. Since these two types of data provided complementary information, merging them increased the overall accuracy of the result.

We also took into account the biological reality that a single cell can give rise to multiple offspring cells, but the reverse is not possible. After this merging process, we arrived at the ultimate relationships of hematopoietic cells in the leukemias (Table S2-Tabel S5).

### NicheNet analysis

NicheNet (version 1.1.1) is a powerful tool for predicting the ligands that drive the transcriptomic changes of target cells. To identify potential ligands that influence cell change, we selected the healthy bone marrow hematopoietic cells and malignant cells as sender cells, respectively, and used the expansion cells as well as the non-expansion cells as receptor cells. To evaluate the top active ligands, we chose the cells that showed expansion as the negative control cells and constructed interactions between ligands and receptors. We listed the top 40 ligands of each cell type and assigned scores to these top ligands corresponding to their ranks.

### pySCENIC analysis

pySCENIC could infer transcription factors, gene regulatory networks and cell types from single-cell RNA-seq data. Activated regulons in each hematopoietic cells were analyzed using pySCENIC with raw count matrix as input. SCENIC UMAP plots could show both highly variable genes and cell type classifications. Also, regulon specificity scores (RSS) across different cell types were calculated through regulon AUC to predict the condition of cells. The regulon AUC scaled score was generated to compare the potential changing condition of the hematopoietic cells across the leukemia.

### TCGA data analysis

The expression data of TCGA were downloaded. Overall survival (OS) from the TCGA Pan-Cancer Clinical Data Resource (TCGA-CDR) and aml_ohsu_2022 datasets were used to analyze patients’ clinical outcomes. The p-value was calculated and showed.

---

**Supplementary Figure 1.**
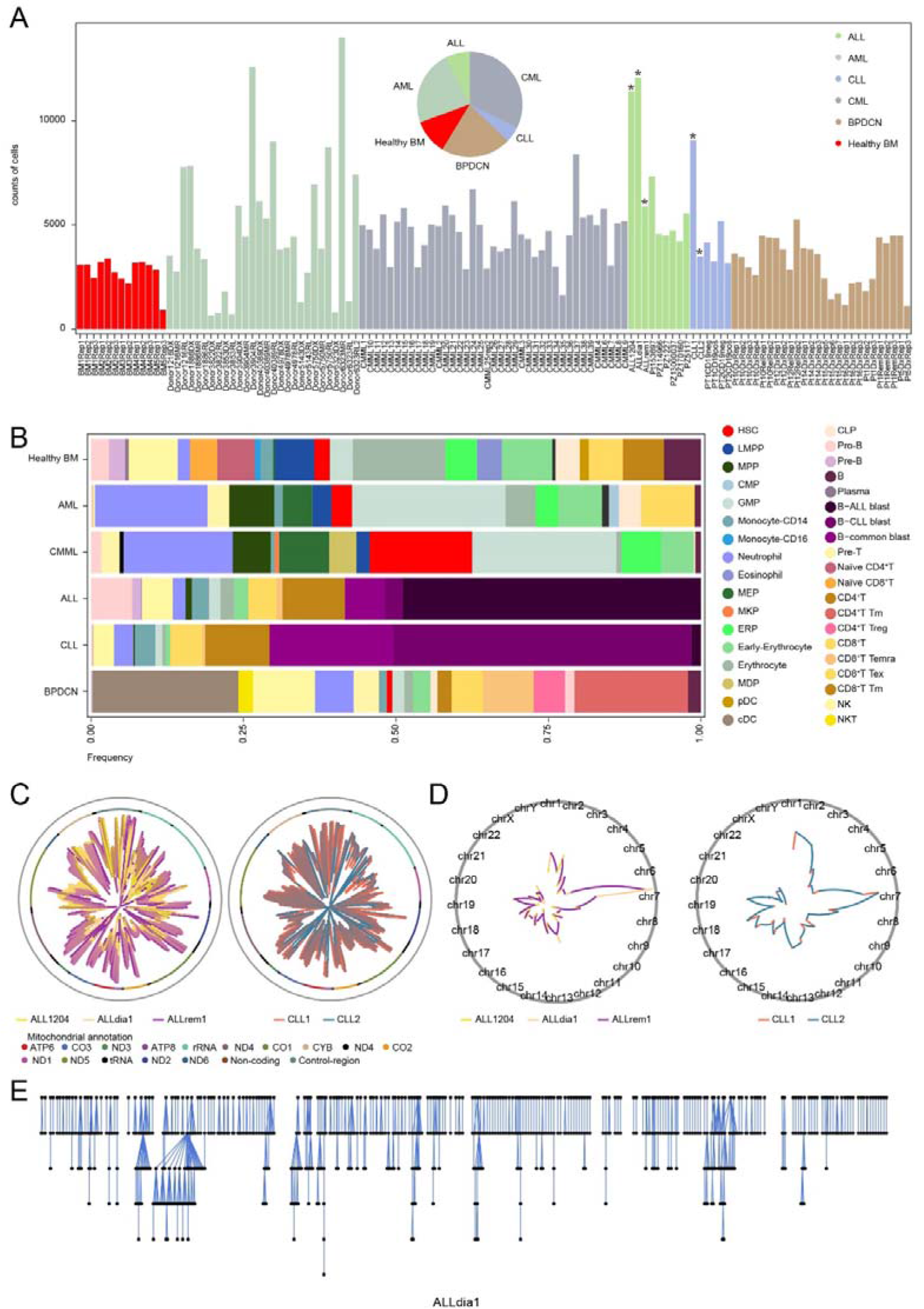
Basic information of the hematopoietic cell subsets in each leukemia type and the coverage and mitochondrial and SNP. (A) Bar plot showing the number of hematopoietic cells collected in each sample. Pie chart showing the proportion of samples from different leukemia types. The newly generated data were labelled. (B) The fraction of hematopoietic cell subsets across leukemia types and healthy bone marrow samples. (C) The coverage of the mitochondrial genome of the samples we newly generated (ALL on the left and CLL on the right) was shown in the inner circle, while the mitochondrial genome was displayed in the middle circle and the outer grey circle represented genome coordinates. The coverage was the sum of single cells. (D) The coverage of chromosome SNPs in the samples we newly generated (ALL on the left and CLL on the right) was shown. The coverage was the sum of single cells. (E) Phylogenetic tree calculated by both mtDNA mutations and SNPs, depicting the cell changing process where ancestor cells transferred to descendant cells in the ALLdia1 sample. Each dot represented a cell.

**Supplementary Figure 2.**
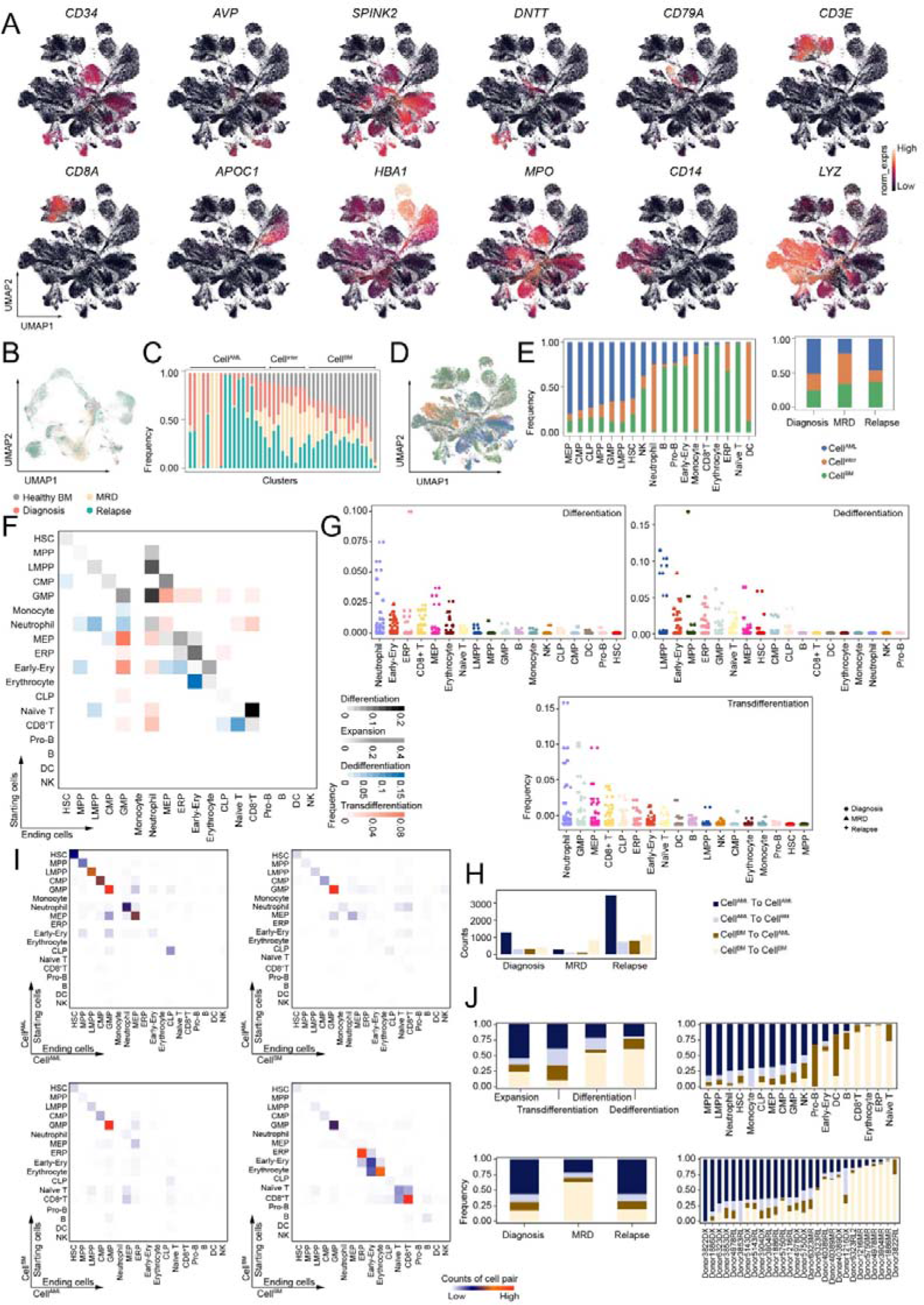
Basic expression information and transcriptional information of AML datasets. (A) UMAP plots showing the marker genes expression for the major lineages of hematopoietic cells in AML datasets. (B) UMAP plot showing the integration of AML and healthy bone marrow scRNA-seq datasets. (C) Bar plot showing the fraction of healthy bone marrow hematopoietic cells in each cluster in (B). The AML cells were classified as cells with a cell state in AML (Cell^AML^), cells with an intermediate cell state (Cell^inter^) and cells with a cell state in healthy bone marrow (Cell^BM^) according to the fraction of healthy bone marrow hematopoietic cells in each cluster. (D) UMAP plot showing the distribution of cells classified in (C) in AML samples. (E) Bar plots displaying the frequency of cells classified in (C) across various cell types and clinical stages. (F) Heatmap showing the frequency of cell changing types in the AML datasets analysis. The cell types in the rows were the starting cells, and the cell types in the columns were the ending cells. Different colors represented different cell changing types. The frequency was calculated through dividing the counts of each cell change type by the total number of all cell changing types. (G) Boxplots illustrating the frequency of cell changing types across the different ending cell types. (H) Bar plot showing the counts of cell changings among cells classified in (C). (I) Heatmap showing the counts of cell changings among cells classified in (C). The cell types in the rows were the starting cells, and the cell types in the columns were the ending cells. (J) Bar plots displaying the fraction of cell changings among cells classified in (C) across the cell changing types, cell types, clinical stages and individual samples, repectively. DX, diagnosis sample; RL, relapse samples; MR, minimal residual disease sample.

**Supplementary Figure 3.**
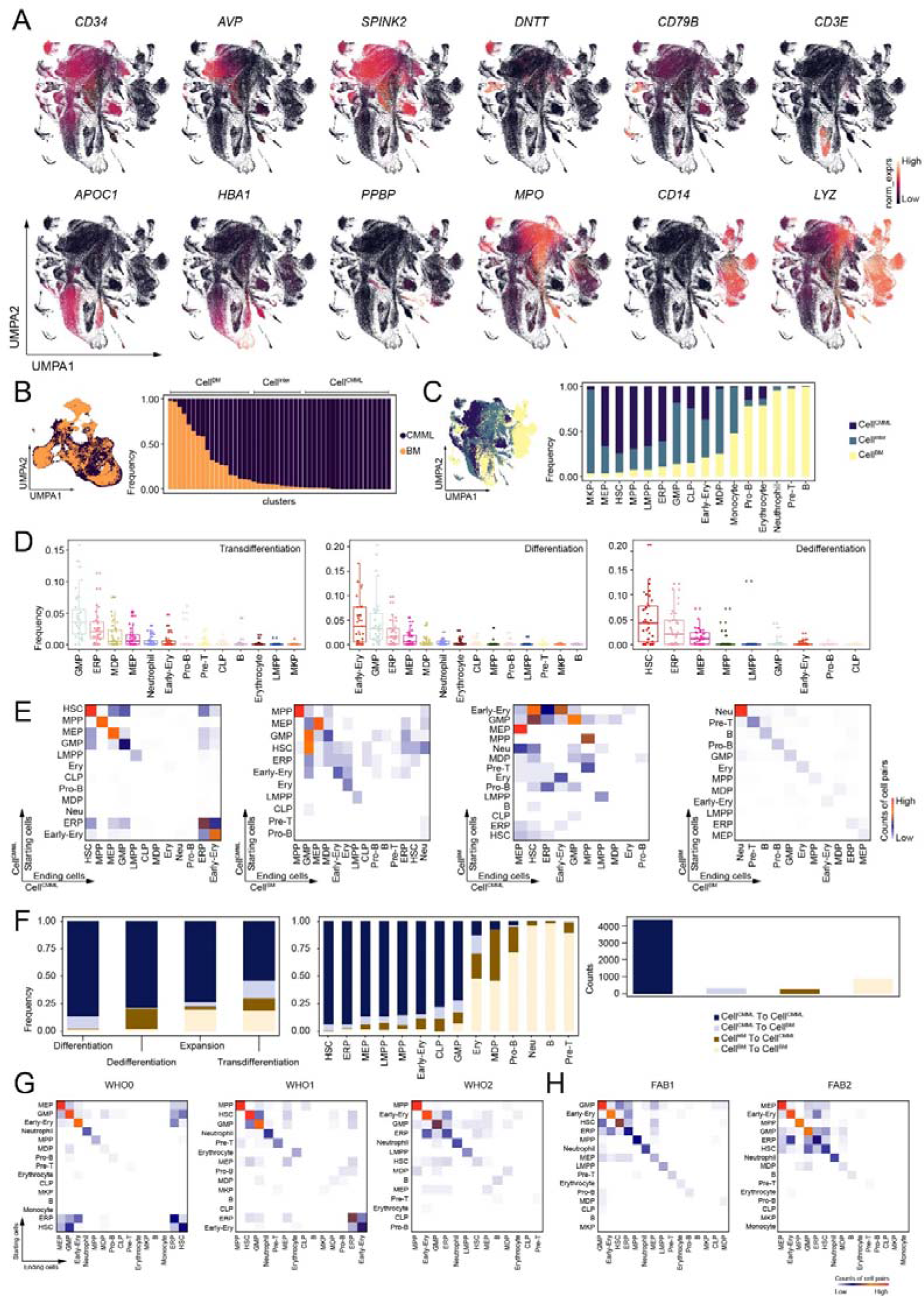
Basic expression information and transcriptional information of CMML datasets. (A) UMAP plots showing the marker genes expression for the major lineages of hematopoietic cells in CMML datasets. (B) UMAP plot showing the integration of CMML and healthy bone marrow scRNA-seq datasets. Bar plot showing the fraction of healthy bone marrow hematopoietic cells in each cluster in. The CMML cells were classified as cells with a cell state in CMML (Cell^CMML^), cells with an intermediate cell state (Cell^inter^) and cells with a cell state in healthy bone marrow (Cell^BM^) according to the fraction of healthy bone marrow hematopoietic cells in each cluster. (C) UMAP plot showing the distribution of cells classified in (B) in CMML samples. Bar plot displaying the frequency of cells classified in (B) across various cell types. (D) Boxplots illustrating the frequency of cell changing types across the different ending cell types. (E) Heatmap showing the counts of cell changings among cells classified in (B). The cell types in the rows were the starting cells, and the cell types in the columns were the ending cells. (F) Bar plots displaying the fraction of cell changings among cells classified in (B) across the cell changing types and cell types (left and middle). Bar plot displaying the counts of cell changings among cells classified in (B) (right). (G) Heatmap showing the counts of cell changings among different cell types across WHO classification. (H) Heatmap showing the counts of cell changings among different cell types across FAB classification.

**Supplementary Figure 4.**
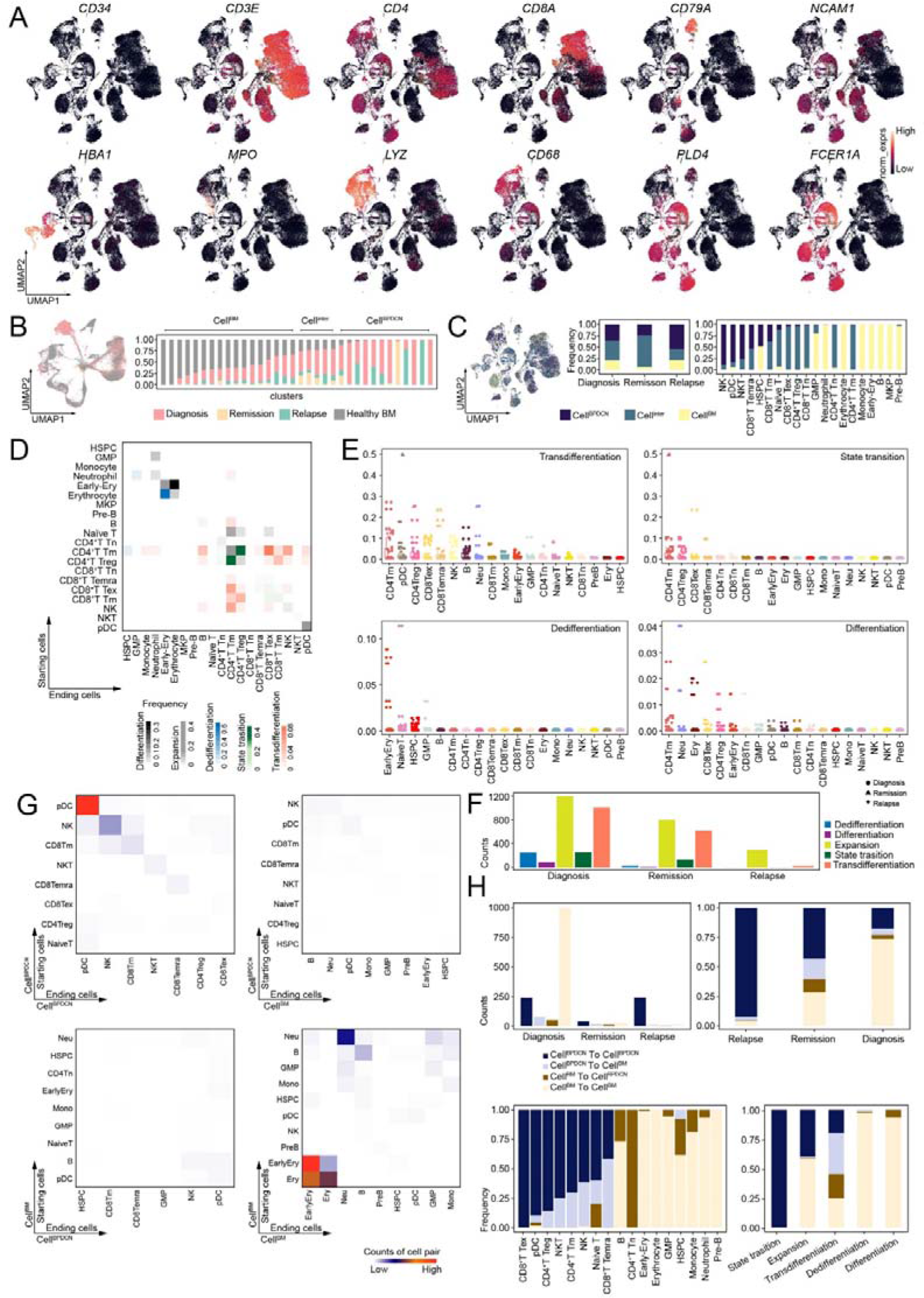
Basic expression information and transcriptional information of BPDCN datasets. (A) UMAP plots showing the marker genes expression for the major lineages of hematopoietic cells in BPDCN datasets. (B) UMAP plot showing the integration of BPDCN and healthy bone marrow scRNA-seq datasets. Bar plot showing the fraction of healthy bone marrow hematopoietic cells in each cluster. The BPDCN cells were classified as cells with a cell state in BPDCN (Cell^BPDCN^), cells with an intermediate cell state (Cell^inter^) and cells with a cell state in healthy bone marrow (Cell^BM^) according to the fraction of healthy bone marrow hematopoietic cells in each cluster. (C) UMAP plot showing the distribution of cells classified in (B) in BPDCN samples. Bar plots displaying the frequency of cells classified in (B) across clinical stages and various cell types. (D) Heatmap showing the frequency of cell changing types in the BPDCN datasets analysis. The cell types in the rows were the starting cells, and the cell types in the columns were the ending cells. Different colors represented different cell changing types. The frequency was calculated through dividing the counts of each cell change type by the total number of all cell changing types. (E) Boxplots illustrating the frequency of cell changing types across the different ending cell types. (F) Bar plot showing the counts of cell changing types across different clinical stages. (G) Heatmap showing the counts of cell changings among cells classified in (B). The cell types in the rows were the starting cells, and the cell types in the columns were the ending cells. (H) Bar plots showing the counts and the fraction of cell changings among cells classified in (B) across the cell changing types, cell types and clinical stages, repectively.

**Supplementary Figure 5.**
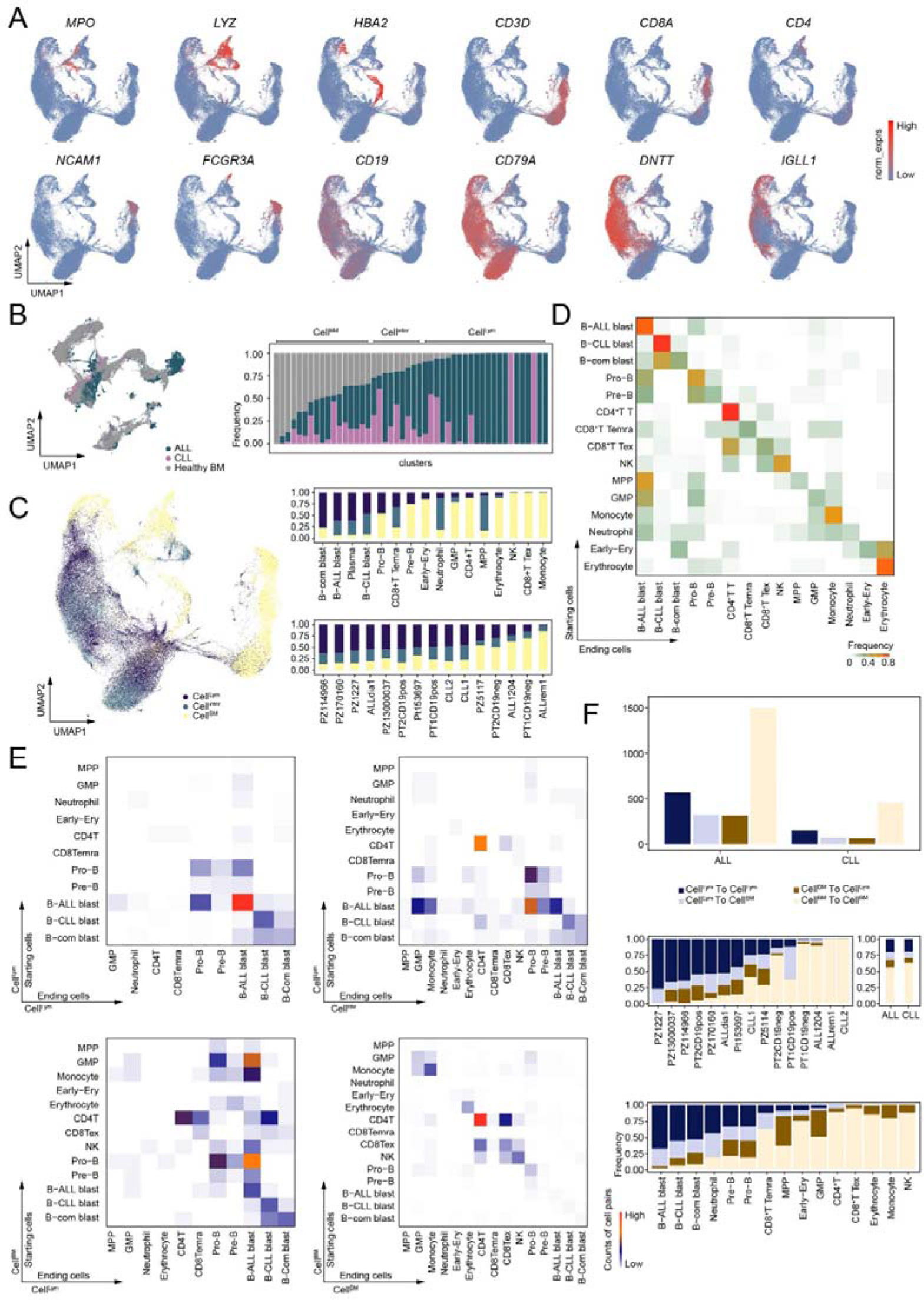
Basic expression information and transcriptional information of lymphoblastic leukemia datasets. (A) UMAP plots showing the marker genes expression for the major lineages of hematopoietic cells in lymphoblastic leukemia datasets. (B) UMAP plot showing the integration of lymphoblastic leukemia and healthy bone marrow scRNA-seq datasets. Bar plot showing the fraction of healthy bone marrow hematopoietic cells in each cluster. The leukemia cells were classified as cells with a cell state in lymphoblastic leukemia (Cell^Lym^), cells with an intermediate cell state (Cell^inter^) and cells with a cell state in healthy bone marrow (Cell^BM^) according to the fraction of healthy bone marrow hematopoietic cells in each cluster. (C) UMAP plot showing the distribution of cells classified in (B) in lymphoblastic leukemia samples. Bar plots displaying the frequency of cells classified in (B) across various cell types and individual samples. (D) Heatmap showing the frequency of cell changing types in the lymphoblastic leukemia datasets analysis. The cell types in the rows were the starting cells, and the cell types in the columns were the ending cells. The frequency was calculated through dividing the counts of each cell change type by the total number of all cell changing types. (E) Heatmap showing the counts of cell changings among cells classified in (B). The cell types in the rows were the starting cells, and the cell types in the columns were the ending cells. (F) Bar plot showing the counts and the fraction of cell changings among cells classified in (B) across the individual samples and cell types, respectively.

**Supplementary Figure 6.**
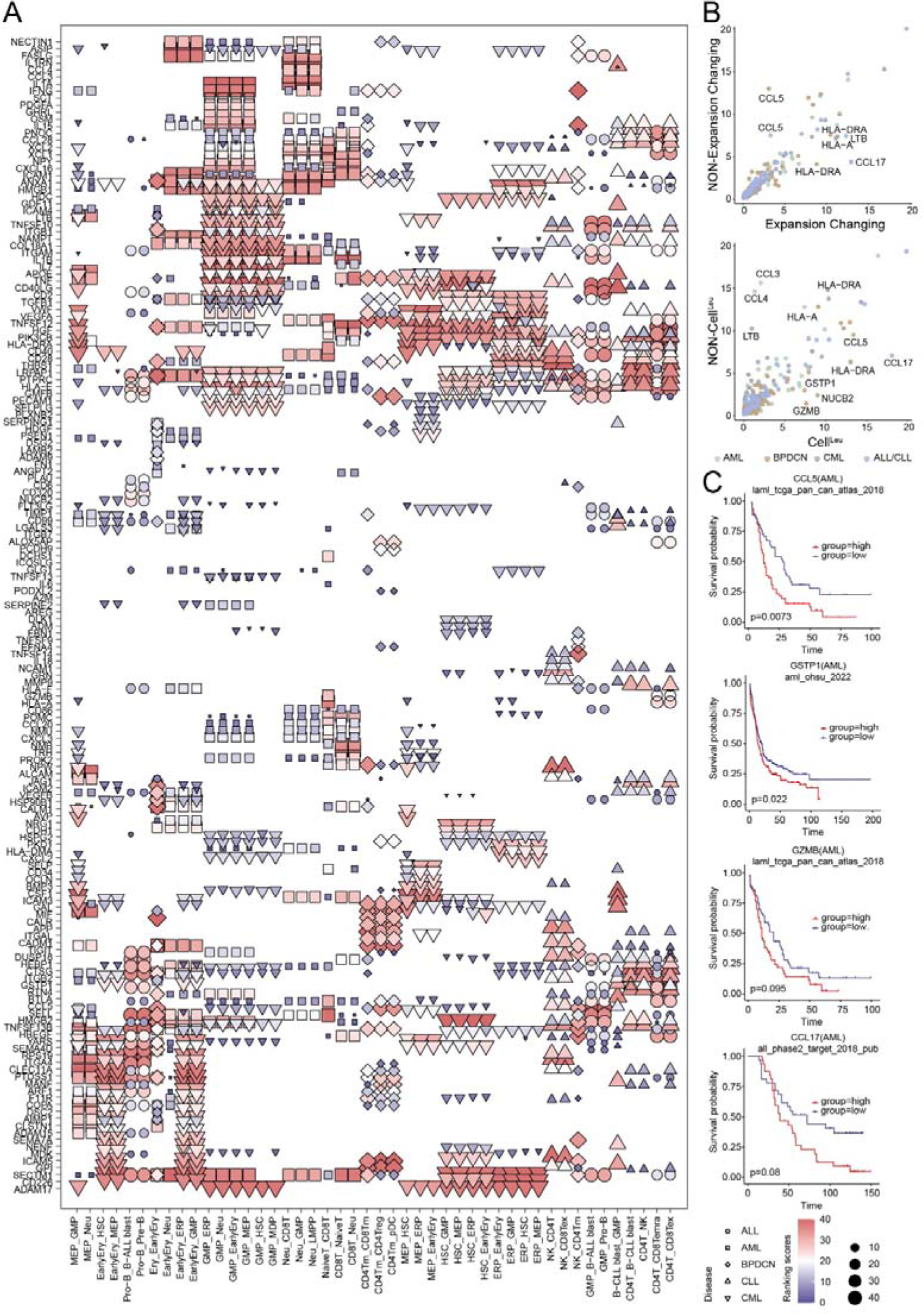
Complexity of the leukemia bone marrow microenvironment. (A) Dot plot illustrating the potential ligands driving the specific cell changings compared with cells underwent expansion across various leukemias within the bone marrow microenvironment. Different patterns represented the different leukemia types. (B) Scatter plot showing the correlation of the potential ligands between the cell changing of expansion and NON-expansion (top), as well as the cells with a cell state in leukemia (Cell^Leu^) and NON-Cell^Leu^ (bottom). (C) Kaplan-Meier plots showing worse clinical outcome in AML patients with the higher expression of CCL5, GSTP, GZMB and CCL17.

**Supplementary Figure 7.**
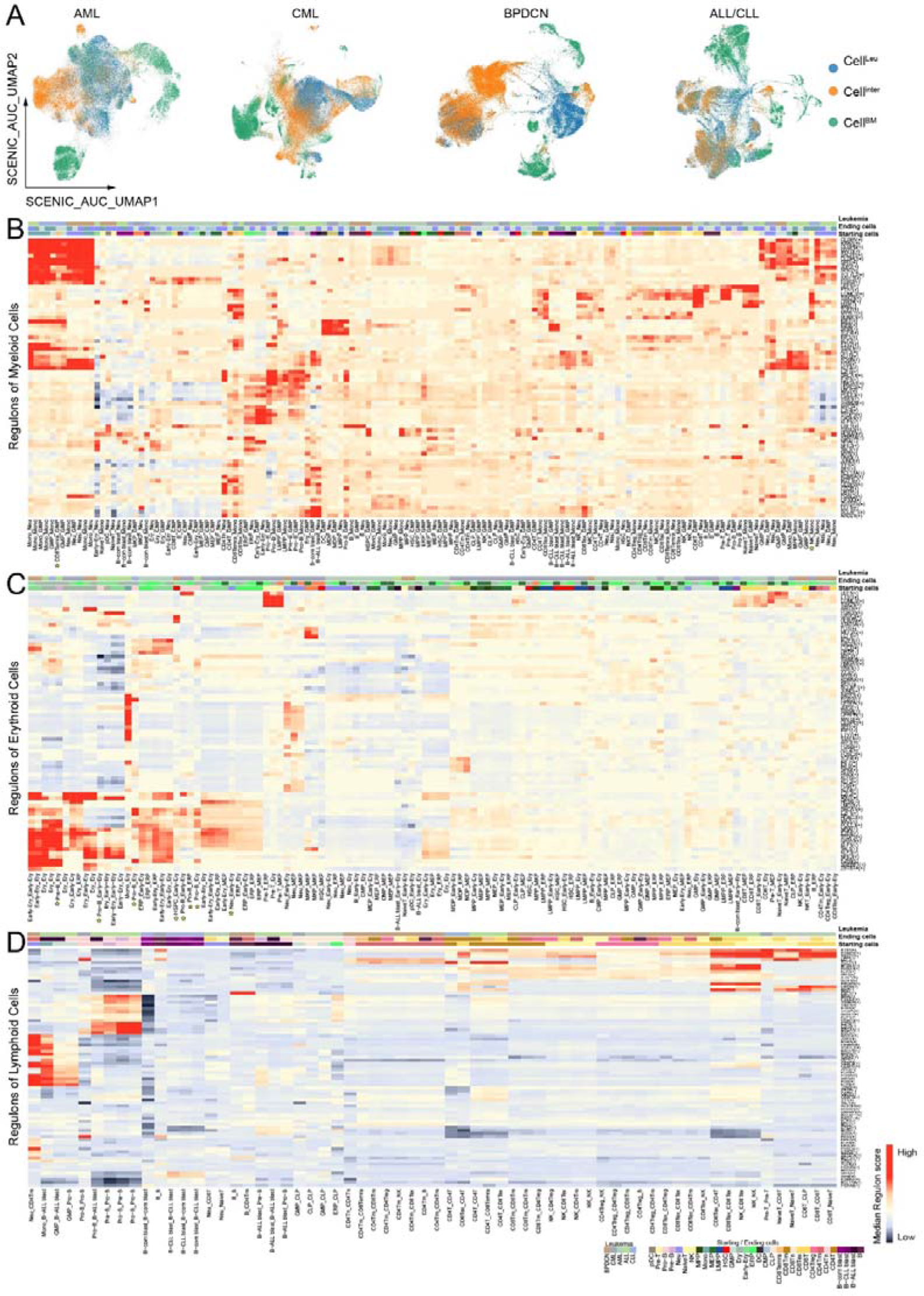
Regulons analysis showed the intermediate cell state during the process of cell changings. (A) SCENIC_AUC_UMAP showing the cells with different cell states across the various leukemia types based on the regulons information. (B) Heatmap showing the regulons of different cell types underwent cell changings to myeloid cells across various leukemias. (C) Heatmap showing the regulons of different cell types underwent cell changings to erythroid cells across various leukemias. (D) Heatmap showing the regulons of different cell types underwent cell changings to lymphoid cells across various leukemias.

**Supplementary Figure 8.**
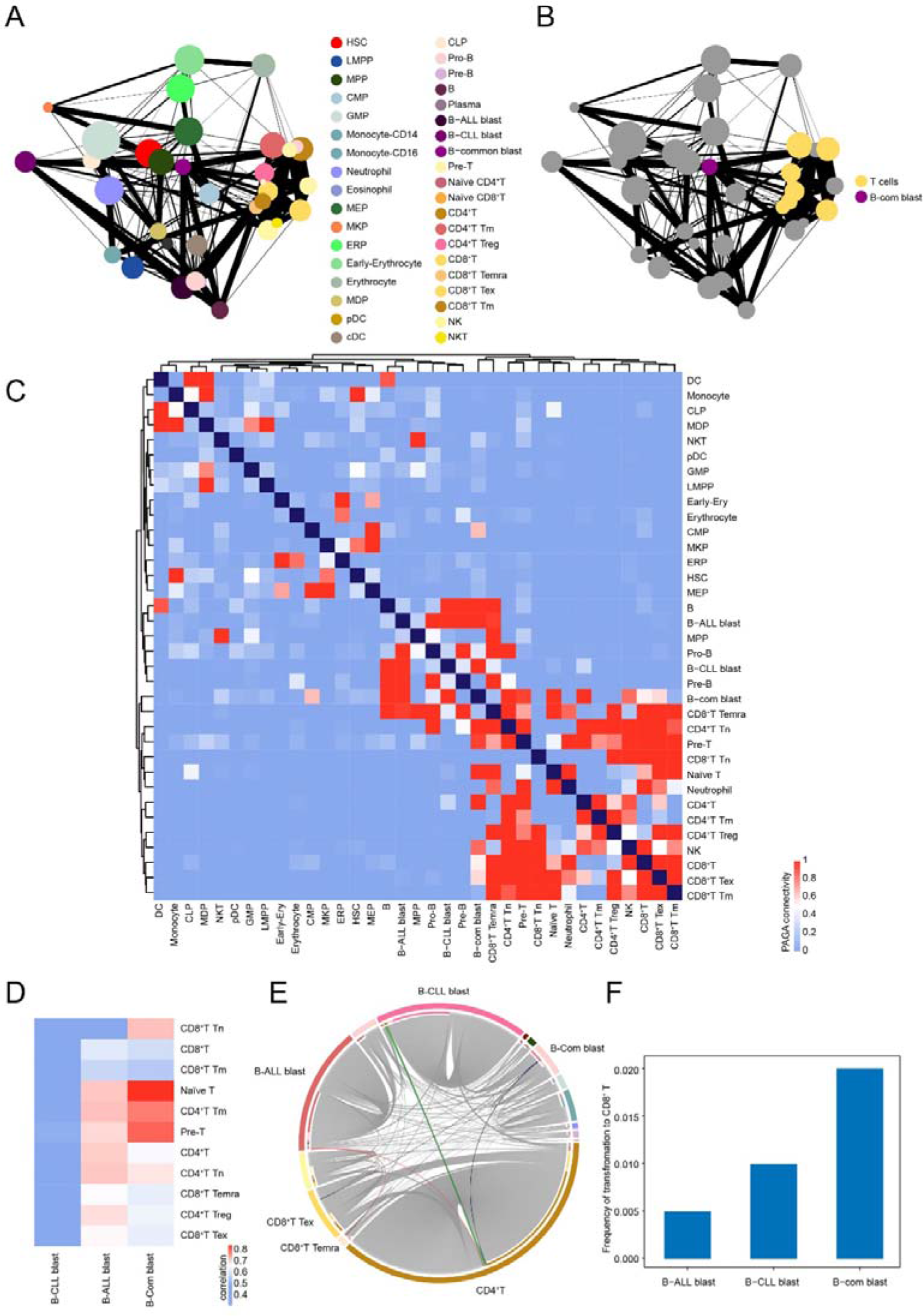
High connectivity between B-common blast cells and T cells. (A) PAGA showing the connectivity of the transferring cells across various leukemia types in this study. (B) PAGA showing the distribution of T cells and B-common blast cells. (C) Heatmap displaying the PAGA connectivity of the transferring cells across various leukemia types. (D) Heatmap showing the correlation of the gene expression between T cell types and B leukemia blast cell types. (E) Circus plot highlighting the cell changings from B leukemia blast cells to T cells. (F) Bar plot showing the frequency of transformation from B leukemia blast cells to CD8^+^ T cells.

## Notes

### Competing Interest Statement

The authors have declared no competing interest.

